# Proteome-based investigation of O-GlcNAcylation in a *C. elegans* model of Ageing and Alzheimer’s disease: Functional Support for Earlier Hypothesis-Generating Findings

**DOI:** 10.1101/2025.09.22.677859

**Authors:** Fernando C. García Olivera

## Abstract

The monosaccharide N-acetyl-glucosamine (GlcNAc), which dynamically modifies serine and threonine residues of nuclear and cytoplasmic proteins, is a key regulator of numerous biological processes. Investigating O-GlcNAc modification in vivo and in vitro remains challenging, making animal models essential for gaining powerful insights into this post-translational modification.

In this study, we conducted a proteomic analysis for identifying O-GlcNAc–modified proteins in both early and adult larval stages of N2 wild-type *Caenorhabditis elegans* and in *aex-3*p::tau(V337M), a nematode model of ageing and Alzheimer’s disease using a high-resolution nano-LC-ESI mass spectrometry approach. We found that O-GlcNAcylation in *C. elegans* is developmentally regulated and is disrupted in a tauopathy model. Wild-type worms transition from RNA processing in larvae to broader regulation of RNA metabolism and protein stability, whereas tau-expressing worms display stress- and signalling-related features. O-GlcNAcylated PDI-2 emerges as a candidate regulator of protein quality control and decrease of O-GlcNAc levels suggests competition with hyperphosphorylated tau. These findings position O-GlcNAcylation as a context-dependent modulator of proteostasis with therapeutic relevance for tauopathy.

## Introduction

According to the World Health Organization, AD is the most prevalent type of dementia, and while age is the strongest known risk factor for its beginning, it is not an unavoidable outcome of ageing. Furthermore, dementia does not only afflict the elderly, up to 9% of cases are young-onset dementia, which is defined as the development of symptoms before the age of 65^1^. However, the molecular mechanisms that mediate the transition from healthy ageing to pathological decline remain incompletely understood.

Among the emerging modulators of neuronal function and proteostasis is O-linked β-N-acetylglucosaminylation (O-GlcNAcylation), a dynamic and reversible post-translational modification (PTM) that regulates thousands of proteins involved in metabolism, stress response, transcription, and proteostasis. Acting as a cellular metabolic sensor, O-GlcNAcylation is tightly coupled to glucose metabolism via the hexosamine biosynthetic pathway (Sun et al., 2016, Di Domenico et al., 2018). In the ageing brain, decreased O-GlcNAcylation has been associated with increased tau hyperphosphorylation and enhanced amyloidogenic processing, both of which are central to AD pathology (Wani et al., 2017; Huang et al., 2020).

O-GlcNAcylation is a post-translational modification involving the addition of a single GlcNAc moiety to serine or threonine residues. It is dynamically regulated by O-GlcNAc transferase (OGT) and O-GlcNAcase (OGA), and although it can occur co-translationally, it is generally considered a post-translational event (Burt et at, 2022). In *Caenorhabditis elegans*, the OGT-1 and OGA-1 enzymes are highly conserved and functionally comparable to their human counterparts. They are essential for maintaining cytoplasmic and nuclear proteostasis. Previous studies have shown that mutations in these enzymes modulate Aβ and tau toxicity in *C. elegans* models, supporting the use of this model organism to explore the molecular roles of O-GlcNAc in ageing and neurodegeneration, as well as to identify small-molecule modulators of O-GlcNAc cycling (Taub et al., 2018; Urso et al. 2020).

Although studies have reported inconsistent findings regarding global protein O-GlcNAcylation levels in AD, there is growing recognition that O-GlcNAcylation is dysregulated in the disease (Wang et al., 2017). However, no clear consensus exists on the specific nature of these alterations, as different studies report divergent results. Some suggest a general decrease in O-GlcNAcylation, while others indicate that changes are more selective or region-specific. Moreover, the interplay between O-GlcNAcylation and other glycosylation pathways may contribute to the complexity of these modifications in AD. For instance, Wang et al. and Frenkel-Pinter et al., (2017) reported significant differences in specific O-GlcNAc-modified peptides in AD brains compared to controls, suggesting that some proteins may undergo reduced modification, whereas others may be upregulated or modified in distinct ways.

While O-GlcNAcylation of other proteins, including the amyloid precursor protein (APP), has been investigated in the context of AD, tau remains a central target due to its critical role in tangle formation. Most of the research has focused on aberrant tau O-GlcNAcylation as a driver of tau pathology in AD, reinforcing the hypothesis that this post-translational modification (PTM) is a crucial modulator of tau function and aggregation (Kielbasa et al., 2024). Nevertheless, given the broad scope of O-GlcNAcylation across many proteins, its dysregulation may affect multiple cellular pathways involved in AD pathogenesis beyond tau.

In this study, we used a proteomic approach the wild-type (WT) N2 strain as a control and a transgenic *C. elegans* expressing the human 4R1N tau isoform carrying the AD-linked V337M mutation, under the control of the pan-neuronal *aex-3* promoter, to systematically identify and profile O-GlcNAcylated proteins associated with ageing and AD pathology across developmental stages. This strategy enabled the identification of stage-specific O-GlcNAcylation patterns under normal conditions and demonstrated that tau expression alters these patterns, particularly in stress and signalling pathways. These results indicate that O-GlcNAcylation modulates RNA metabolism, protein homeostasis, and suggest a mechanistic basis for exploring its therapeutic potential in ageing and neurodegenerative diseases.

## Materials and Methods

### Materials

Agar (Roth Cat. No. 8503); Peptone (Merck Cat. No. 91079-38-8); CaCl2 (Merck Cat. No. 102378); Cholesterol (Sigma-Aldrich Cat. No. C8503); CuSO_4_. 5H_2_O (Sigma-Aldrich Cat. No. C8027); NaCl (Sigma-Aldrich Cat. No. S7653); K_2_HPO_4_ (Merck Cat. No. P290); KHCO_3_ (Sigma Aldrich Cat. No. 237205); KCl (Sigma-Aldrich Cat. No. 9541); KH_2_PO_4_ (Roth Cat. No. 3904.1); Na_2_HPO_4_ (Sigma-Aldrich Cat. No. S7907); MgSO_4_ (Roth Cat. No. 0261.1); NaOH (Sigma-Aldrich Cat. No. 1310-73-2); NaOCl (Roth Cat. No. 9062-3); Tris (Tris-(hydroxymethyl)-aminomethan, Roth Cat. No. 4855.2); Triton X100 (Cat. No. 9002-93-1); DTT (Cat. No. D0632); NH_4_CO_3_ (Cat. No. 1066-33-7); (Succinylated *Titricum vulgaris* Lectin sWGA (MagneZoom™, GlycoMatrix cat. No. 20120079-1); Binding Buffer (GlycoMatrix cat. No. 21511271-1); Elution Buffer (GlycoMatrix cat. No. 21511313-1).

## Methods

### Worms Synchronization

WT N2 (Bristol) and the *aex-3*p::tau(V337M) strains (University of Washington, Seattle, USA) were used for our experiments. The *aex-3* promoter drives the expression of the human tau V337M transgene predominantly in neurons and is commonly used in *C. elegans* for pan-neuronal expression, making it a suitable genetic tool for modeling tauopathy and investigating neuron-specific effects of post-translational modifications (Pir et al., 2019). A solid sterile Nematode Growth Medium (NGM) was prepared and sown with Escherichia Coli OP50 (Protein analytics, Giessen (HE), Germany) for feeding on 94 x16 mm plates (Greiner Bio-One, Frickenhausen, Germany) and was used to propagate *C. elegans* strains according to standard procedures (Hibshman et al, 2021). All strains were grown at a temperature of 20°C. These plates were grown for 24h in an incubator (MAGV, Rabenau, Germany). Every two to three days, they were moved to a plate containing a new NGM medium. Six plates of adult worms of each strain, to obtain ∼ 30.000 eggs, were bleached using the alkaline hypochlorite method to prepare synchronized animals, respectively (Hibshman et al, 2021). In a nutshell, worms were removed from culture plates using M9 buffer (22 mM KH_2_PO_4_, 42 mM Na_2_HPO_4_, 86 mM NaCl and 1 mM MgSO_4_). After removing all the bacteria from the supernatant, they were lysed in a solution of alkaline hypochlorite for 1 minute and neutralized by using M9 medium; this process was monitored under a dissecting microscope and repeated six times until worms broke apart. Debris was washed with M9 by centrifugation at 2000 rpm for 3 min at 4°C (Hettich® Universal 32R Centrifuge, Tuttlingen, Germany) until the solution was cleared (eight wash cycles). The obtained eggs were then cleaned by centrifugation at 1800 rpm for 3 min using M9 buffer at 4°C and allowed to nutate overnight before hatching in 2 mL of M9 in a climatic chamber (Liebherr®, Ochsenhausen, Germany) at 20°C.

### Worm lysis and protein extraction

Aliquots of worm eggs of N2 and *aex-3*p::tau(V337M) strains were pipetted in 15mL tubes containing 1mL M9 medium. These were let mature in the mentioned solution overnight. Aliquots of L1 hatchlings were seeded onto brand-new NGM plates containing OP50 culture. this population was grown at 20°C for 10 days. Another L1 *C. elegans* larvae of corresponding wild and transgenic strains were rinsed three times and kept for protein extraction.

On day 11, worms were collected by adding 3 mL of M9 buffer to each plate. The plates were gently swirled, and the solution was filtered through a 50 µm mesh (Cytecs, Muenster, Germany) and thoroughly rinsed with M9 buffer. This washing procedure was performed twice to obtain a synchronized population of one-day-old and middle-aged adult L4 stages. Nematodes remained on the 50 µm mesh were carefully transferred into a 15mL tube and centrifuged for three minutes at 1800rpm at 4 °C. The supernatant was removed, and the worms were resuspended with 10 mL of M9 buffer and centrifuged three times discarding the supernatants. The washed L1 and L4 populations were then transferred to 1.5-mL microfuge tubes separately and concentrated by centrifugation (1800rpm, 3 min, 4 °C), the supernatant was removed, and 300 uL of ice-cold lysis buffer was added (150 mM NaCl, 20 mM Tris pH 7.4, 1.0% Triton X-100) supplemented with protease inhibitor cocktail. The suspended worms were pelletized again at 1800 x rpm for 3 min and frozen the tube in liquid nitrogen followed immediately by quickly thawing in a 37°C water-bath and disrupted by subjecting the samples to vortex pulses cycles of 15s each (Vortexer, Heidolph, Schwabach, Germany). This freeze-thaw cycle was repeated three times with cooling on ice between pulses. After the second freeze-thaw cycle, samples lysis was enhanced by using two ultrasound pulse cycles of 15s (Ultrasonic bath, Elmasonic S40, Singen, Germany). Lysates were maintained on ice and examined using a light microscope to verify successful lysis of the worm.

The sonicated worm lysates were centrifuged at 14,000g for 10 min at 4°C to pellet any worm-debris. Subsequently, the lysates supernatants were transferred to a 15 mL conical tube at 20°C–25°C. Worm lysates were made up the total volume to 1,8 mL. Total protein amount was measured in the range of 1–4 mg per tube.

### Protein precipitation, resuspension, and quantification

3 volumes (6 mL) of ice-cold acetone were added to the worm lysates to precipitate proteins and mixed well by inverting a few times. The samples were incubated overnight at −20°C to aid the precipitation. Precipitated proteins were pelleted by centrifugation at 4,000 g for 30 min at 4°C. The supernatants were decanted carefully, without disturbing the pellets. Any remaining acetone was removed, and then air dry the pellet for 15 min by leaving the uncapped tube at 20°C–25°C.

Pellets were resuspended in 200 μL of a buffer containing 100 mM NH_4_HCO_3_, by pipetting up and down. Protein quantification was performed using the Pierce BCA protein assay kit (Cat. No. A55864) and measured with an absorbance microplate reader (BioTek ELx800™, Washington, USA) as per manufacturer instructions. The protein lysates were reduced for 30 min at 37°C with 5mM DDT and subsequently alkylated using 15mM iodoacetamide for 30 min at room temperature in the dark and digested overnight at room temperature with sequencing grade trypsin (MS gold-grade, Mannheim, Germany) at an enzyme ratio of 1:50 (w/w) to a final reaction volume of 500 μL. Samples digestion was stopped with trifluoroacetic acid at a final concentration of 0.1%.

To remove any remaining insoluble material that had not been digested, samples were centrifuged at 14,000 g for 20 min at room temperature. The supernatant, which included the peptide mixture, was then collected in Lobind® tubes (Eppendorf cat. No. 56251, Hamburg Germany.

The insoluble particle’s remaining peptides were extracted by adding 0.5 mL of 0.1% TFA in water, pipetting the pellets back into suspension, and then centrifuging them once more for 20 min. The supernatant was combined with the prior one, desalted by using strong C18 solid-phase extraction cartridges (Macherey-Nagel cat. No. 730011, Dueren, Germany). Detergent traces were removed by incubation and centrifugation using Pierce™ removal spin columns (Thermo Fisher Scientific Cat. No. 87777, Dreieich, Germany), and dried using SpeedVac (Eppendorf, Concentrator Plus™, Hamburg, Germany) completely.

### Peptide enrichment

We employed a label-free alternative to enrich and detect O-GlcNAc-modified proteins, by employing succinylated wheat germ agglutinin (sWGA), a lectin that binds specifically to terminal N-acetylglucosamine (GlcNAc) residues. Succinylation of WGA reduces its affinity for sialic acid and other non-specific glycan structures, thereby increasing its selectivity toward β-linked GlcNAc residues typical of O-GlcNAc modifications on intracellular proteins (Kupferschmid *et al*., 2017; Dupas et al., 2022,). Unlike native WGA, which exhibits broader carbohydrate-binding activity, sWGA offers improved specificity for cytoplasmic and nuclear O-GlcNAcylation without substantial cross-reactivity to complex N-glycans or membrane-associated glycoconjugates (Sečová et al., 2024, Moniaux et al., 2025). This increased specificity makes sWGA particularly suitable for applications such as lectin blotting, affinity purification, and gel-based mobility shift assays targeting O-GlcNAc-modified proteins (Kubota et al., 2017; Tang et al., 2024; Zhang et al 2024). Thus, sWGA provides a reliable and widely accepted tool for detecting O-GlcNAc in diverse biological samples, especially when antibody-based approaches may lack sensitivity or cross-react with other glycans.

For our study, 1mg of dried peptide pellet were dissolved in 1 mL of a buffer 50mM sodium phosphate, 7.0, containing 20mM NaCl (Binding buffer) and then incubated overnight at 4°C with a 200 µL slurry containing *s-WGA –* MagneZoom*™* (Dublin Ohio, USA) beads using rotational mixing (IKA Loopster basic, Staufen, Germany). The beads were precipitated by centrifugation at 1000 g for 5 min and washed by adding 0.5mL of Binding/Wash Buffer and centrifuging again to remove the supernatant, the wash process was repeated two additional times.

Tubes containing samples was removed from the centrifuge. Bound peptides were eluted by adding 300 µL of 50mM Sodium Phosphate buffer containing 0.1-0.2M N-acetylglucosamine and 500mM NaCl (Elution buffer). The beads were resuspended and incubated at room temperature for 15 min with rotational mixing. Following incubation, the beads were centrifuged at 1000 g and the supernatant was transferred to a clean tube. The incubation and elution were repeated three times using elution buffer.

The flow through was then collected and desalted using strong C18 cartridges solid-phase (Chromabond) and dried using a SpeedVac concentrator for the MS analysis.

### Nano-LC-MS/MS analysis and evaluation of O-glycan

1µg from the Glycopeptides enriched above were redissolved in 0.1% formic acid and loaded onto a 50 cm µPAC™ C18 column (Pharma Fluidics, Gent, Belgium). The peptides were eluted using a linear gradient from 9 to 50% of 20 mM NH_4_OCH_3,_ pH 10, in 90% acetonitrile over 240 min at a constant flow rate of 300 nL/min (Thermo Fisher Scientific™ UltiMate™ 3000 RSLC nano), infused via an Advion TriVersa NanoMate (Advion BioSciences, Inc. New York, USA) and analysed by an Orbitrap Eclipse Tribrid mass spectrometer (Thermo Fisher Scientific, Waltham, MA, USA. Each precursor ion (charged 2-7+) was separated in the quadrupole (selection window: 1.6 m/z; dynamic exclusion window set to 30 s and a 3 s scan cycle by using EThcD fragmentation (Maximum Injection Time: 250 ms; normalized Collision Energy: 25%) and a resolution of 30,000; AGC; 5.0e4) to trigger HexNAc oxonium ion fragments.

Raw mass-spectra data files were processed for intact O-glycopeptides identification using Mascot Distiller (Matrix Science, London, UK, v2.8.5.1 version), a software that process high-quality peak lists for Mascot database searches. The following parameters were set; Species, *C. elegans*; enzyme specificities (Trypsin, Digest C Term (KR) with a maximum of two missed cleavages; fixed modification, carbamidomethylation of Cys (C); variable modifications were HexNAc (ST) and Methionine oxidation (M). The mass accuracy was 0.08 Da for-precursor ions and 0.05 Da for the fragment ions. Annotation results were filtered to obtain peptide spectrum matches (PSMs) with a ≤1% FDR (False Discovery Rate). The search of corresponding O-glycopeptides was conducted against the current secretome UniProtKB/Swiss-Prot database, contaminants, and UP1940 for *C. elegans*.

Peptides among this range were considered O-GlcNAcylated candidates and were manually inspected by diagnostic of HexNAc oxonium ions (m/z 204.0867 with 138.055 and/or 144.066), observation of a ∼203.0794 Da neutral loss, and sequence support from b/y ions of the modified residue. Expectation Value (E-Value) less than 0.05 and Mascot ion score more than 20 and <30 were set as acceptance criteria. HexNAc on S/T with an ion score ≥30, were considered as strong evidence of O-GlcNAc glycopeptide after positive manual validation with a mass error between 5 - 10 parts per million (ppm) and scores of 40 or higher were designated very high-confidence (Hensbergen *et al*., 2022; Veith et al., 2023). Half-box plots representation of distribution between parameters for all the identified sequences was performed using OriginPro 2025 (Origin Lab Corporation)

### Identification of Orthologs

Orthologs protein identification was performed using UniProt database in *Homo sapiens* sequence identity of 100% and OrthoList2, a database that compiles *C. elegans* and human orthologs based on a 2018 meta-analysis. (Table S1; see Supplementary Information), according to conservative standards commonly applied in recent large-scale studies of protein orthology and comparative genomics (Kim *et al*., 2018; Ahmad *et al*., 2025).

### Protein-Protein Interaction Network (PPI)

A protein-protein interaction network was constructed using the STRING v12.0 database to identify molecular associations. For the N2 strain, a maximum of 21 interactions per protein was permitted. For proteins detected in aex-3p::tau(V337M), only one interaction per protein was allowed, following established methodological guidelines for this type of analysis (Papik et al., 2025). A high confidence level (0.7) was chosen as the confidence limit. Protein-protein interactions were predicted using text mining, experiments, neighbourhood, occurrence, co-expression, and information imported from publicly available databases of the full STRING network (Szklarczyk *et al*., 2023).

## Results

### Global Identification of O-GlcNacylated proteins

We performed a proteomic identification of O-GlcNAcylated targets to examine changes in protein O-GlcNAcylation as nematodes age, and changes during the progression of AD pathology. The characterization phase initiated by using collected samples of N2 WT and *aex-3*p::tau(V337M) *C. elegans* to search for sequences containing motifs corresponding to O-GlcNAcylated proteins.

O-GlcNAcylated peptides annotated by Mascot were subjected to additional layers of validation to ensure confident assignment. Only PSMs with scores exceeding the identity threshold (expect <0.05) were considered further. Inspection of the corresponding MS/MS spectra revealed diagnostic HexNAc oxonium ions at serine/threonine residues (m/z 204.0867 and related fragments), as well as neutral-loss signals (−203 Da) characteristic of O-GlcNAc. Oxidation (M) was also reported on one peptide as variable PTM. Backbone fragmentation was assessed to determine coverage of the modified residue. The analysis included both intact b and y ions as well as neutral-loss derivatives, specifically y* ions with loss of ammonia (−NH_3_) and y ions with loss of water (−H_2_O). Only ions with a mass error of 5-10 ppm or less were considered. Both singly and doubly charged states were evaluated using extracted ion annotations for each strain (please refer to Supplementary dataset 1). Peptides with Mascot ion scores greater than 20 and <30 were considered candidate O-GlcNAcylated peptides. Scores above 30 indicated high confidence and scores of 40 or higher represented excellent confidence.

### High-Confidence Peptides (Score ≥30)

Most annotated O-Glycopeptides achieved scores above the significance threshold, ensuring reliable identification. Representative examples included:

PNSRHDNVSPSK (score 41.0; Mr(calc): 1742.81), identified with HexNAc modifications at S9 and S11. LLVADILACNDDTPASAMMAGNGPVATMSLQVK (score 43.2; Mr(calc): 3519.70), containing a glycosylation at S29. HGGTTRTADAIRYATK (score 33.6; Mr(calc): 2124.04), observed in multiple HexNAc-modified variants in N2 L4 and *aex-3*p::tau(V337M) L1.

These high-scoring peptides demonstrated extensive fragment ion coverage and reproducible detection across replicate (n=2) runs. Collectively, they provided confident assignments to proteins such as glutamic acid–rich protein, K-homology protein domain, and VWFA-domain containing protein.

### Candidate Peptides (Score <30)

A smaller subset of annotated peptides did not reach the Mascot confidence threshold; however, these O-glycopeptides were classified as reliable identifications based on consistent tandem mass spectrometry (MS/MS) fragmentation patterns. These included: VGLIAARRTGR (score 22; Mr(calc): 1371.79), mapping to ribosomal protein uL2 (RL-8_CAEEL). RARDSASSSSSHSK (score 23.0; Mr(calc): 2477.10), derived from a spliceosome-associated protein CWC15_CAEEL. DELPAIRLISLEEDMTK (score 22.2; Mr(calc): 2378.18), exhibiting HexNAc ions on both Ser and Thr. MFITRGLILISLLFVFVMTDDTHDK (score 22.1; Mr(calc): 3346.71), heavily modified with HexNAc and oxidation. TFDFRADKILESLTNSLK (score 22.2; Mr(calc): 2300.19), derived from a NADP-dependent oxidoreductase domain-containing protein.

While these sequences did not surpass the significance threshold, their pExpect values were below 0.05. Such candidates require further validation using targeted MS/MS or synthetic peptide standards. Table S1 and supplementary dataset 2 present a comprehensive summary of all the annotated Mascot sequences.

Collectively, peptide data were processed and evaluated to obtain proteomics profiles for WT N2 and *aex-3*p::tau(V337M). The monoisotopic mass distribution exhibited an asymmetric pattern, with pronounced concentration in medium and high-mass range with observation of greater O-GlcNAc-modified peptide diversity in the WT strain (Fig. 1A). Ion score values (Fig. 1B) were also more variable in N2, with higher scores than *aex-3* strain, reflecting more confident peptide identifications. Peptide length distributions (Fig. 1C) followed a similar trend: N2 O-GlcNAc-modified covered a broader range of lengths, whereas *aex-3* group peptides were predominantly shorter and less variable. These results indicate that N2 samples displayed greater heterogeneity in mass, score, and length compared to *aex-3*, supporting consistent separation of identification parameters between groups.

**Figure 1.**
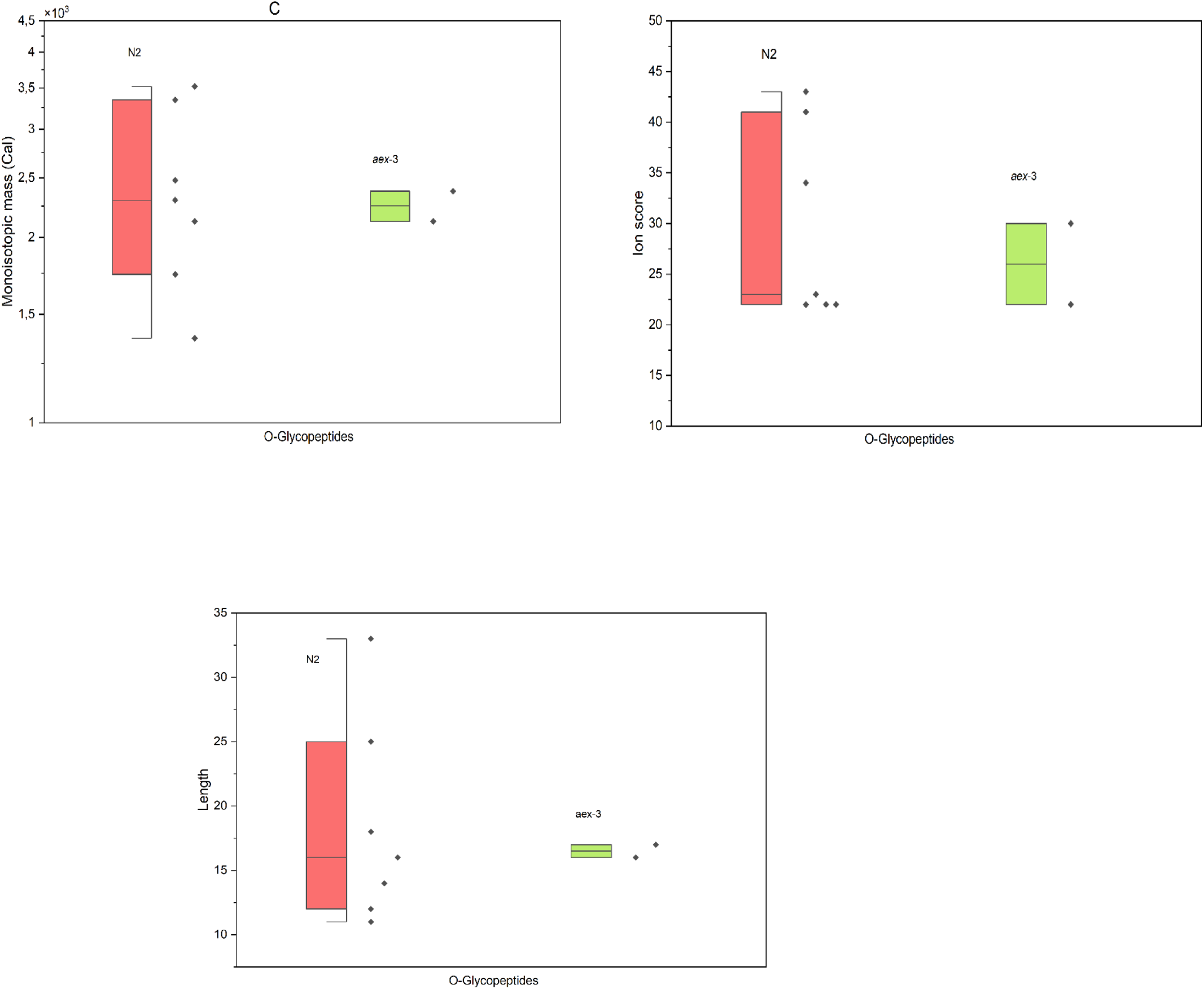
Half-box plots showing the distribution of peptide identification parameters between WT N2 (peach) and *aex-3*p::tau(V337M) (green) (n=2). (A) Monoisotopic molecular weight (Ma Cal), (B) Ion score, and (C) Peptide length. Each plot displays the distribution density, individual peptide values, and the mean ± error bars for each condition.

### Integration of O-GlcNAcylation with Protein Networks and Pathways in Ageing and AD model

Functional enrichment analysis of O-GlcNAcylated proteins in the N2 strain was conducted using STRING interaction network (Fig. 2A). In the BP category, gene ontology (GO) annotations indicated significant enrichment in mRNA splicing, RNA processing (GO:0000398, FDR: <0.01), mRNA processing (GO:0006397, FDR:<0.01), and gene expression (GO:0010467, FDR: < 0.05) as shown in Fig. 3A. The MF category was enriched for RNA binding (GO:0003723, FDR: < 0.05) (Fig.3B). For the CC category, the enriched genes were primarily associated with spliceosomal complex (GO:0005681, FDR: <0.05) as well as ribonucleoprotein complex (GO:1990904, FDR: <0.05) (Fig. 3C). Reactome annotations indicated significant involvement in mRNA splicing-Major Pathway, mRNA splicing, processing of capped intron-containing Pre-mRNA and metabolism of RNA (FDR: < 0.0001) (Fig. 3D). Interestingly, InterPro domain enrichment identified a significant overrepresentation of K homology (KH) domains in the dataset. The KH domain, KH domain type 1, and KH domain superfamily were all enriched (FDR: <0.001) (Fig. 3E and supplemental dataset 3).

**Figure 2.**
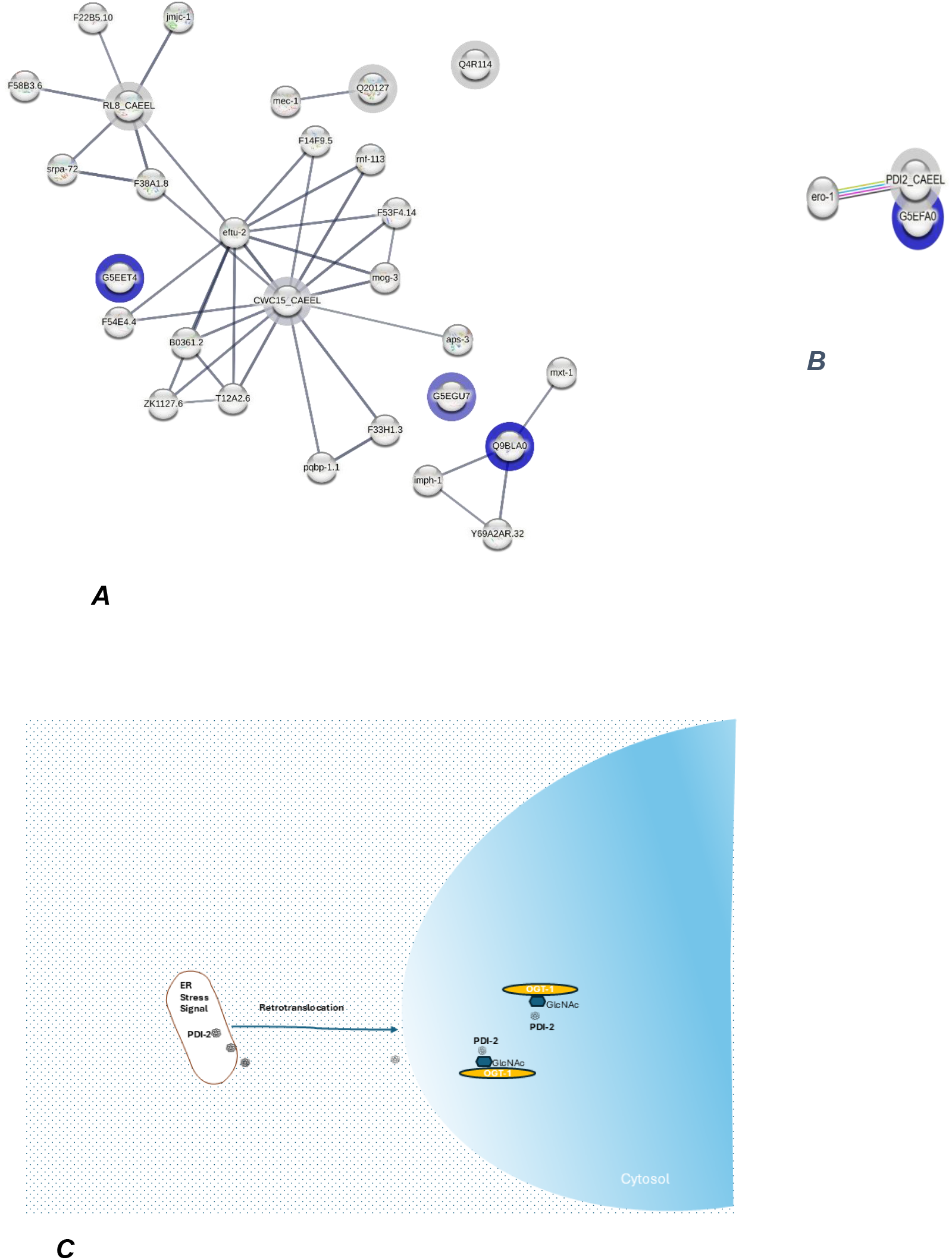
Protein-protein interaction networks from O-GlcNAcylated proteins in young and adult A) WT N2, B) *aex*3p::tau(V337M). Proteins identified with a blue halo represent those with high and very high confidence (ion score ≥30 and pExpect: 0.05). C) Non-canonical O-GlcNAcylation: Relocated PDI-2 modified in cytosol.

**Figure 3.**
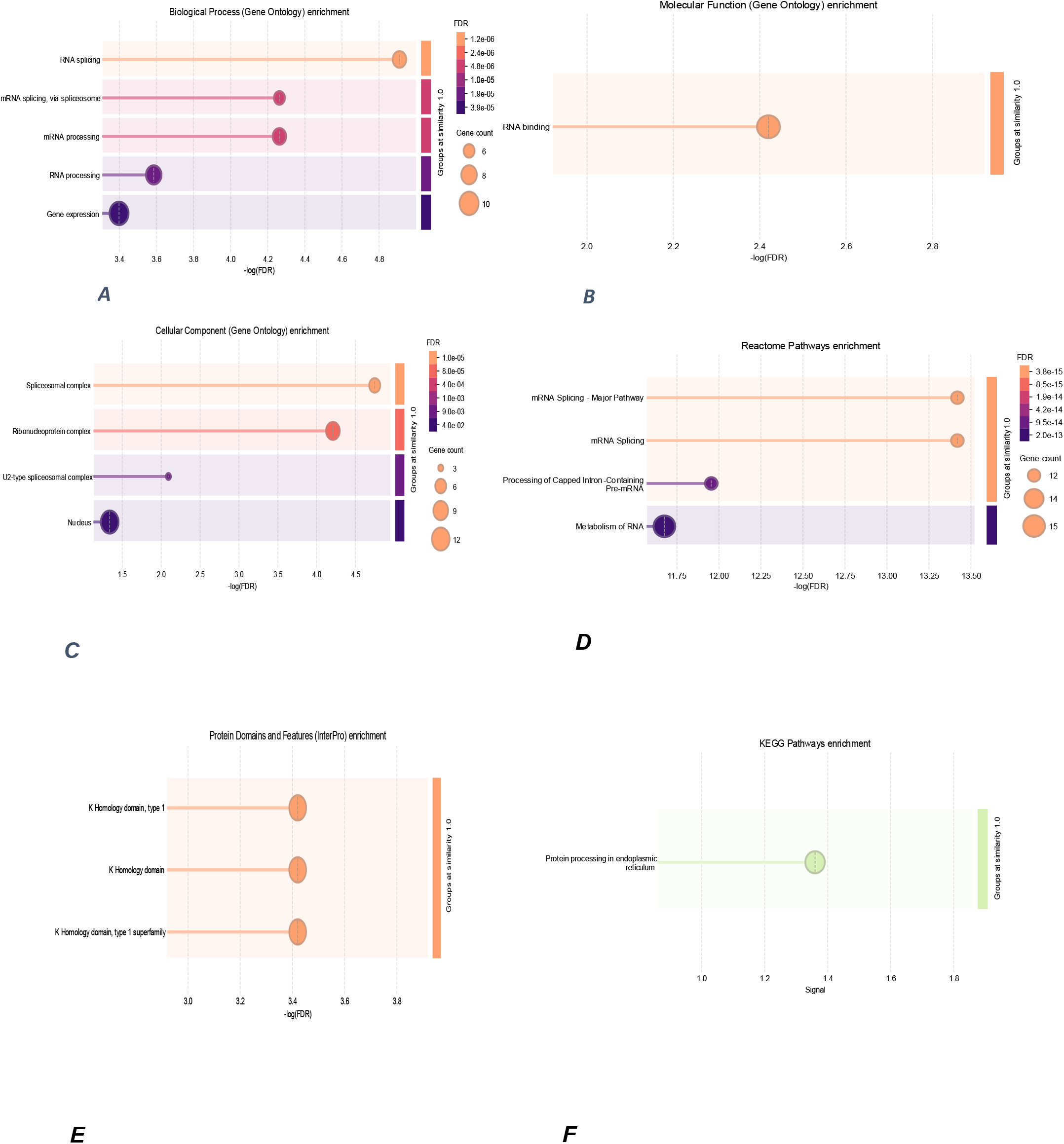
A) Biological process, B) Molecular function, C) Cellular component, D) Reactome Pathways, E) Protein Domains and features enrichment in the WT N2 strain, F) KEGG Pathways enrichment in the *aex-3* transgenic strain.

In parallel, PPI enrichment was evaluated in key signalling pathways of O-GlcNAcylated proteins in *aex-3*p::tau(V337M) strains (Fig. 2B). While this network did not produce any significant interaction in biological processes or molecular functions at L1 and L4 life points, subsequent KEGG pathway analysis identified a significant enrichment in protein processing within the endoplasmic reticulum (ER) (cel04141, FDR: < 0.0001) (Fig. 3F) as well as in WBP annotations associated with the unfolded protein response variant (WBPhenotype:0001719, FDR: <0.05) at adult L4 stage (supplemental dataset 3).

To investigate conservative homology associated with ageing and AD pathology in humans, we performed a query of *C. elegans* O-GlcNAcylated proteins against Ortholist 2 (Kim et al., 2018) and UniProt Database, a percentage of 78% of the queried genes were identified to have human orthologs (Table S1). Comparison of ortholog identification revealed that seven genes were annotated in UniProt as having human orthologs, whereas OrthoList2 (Table S2) identified only five. Moreover, two additional genes were found to lack orthologs in both databases. This discrepancy reflects differences in database coverage and curation criteria. Nonetheless, this integrative approach highlights evolutionarily conserved molecular targets between humans and the *C. elegans* AD model, providing a basis for further interpretation in the context of ageing and AD mechanisms.

## Discussion

Our proteomic profiling revealed stage- and strain-specific protein expression patterns in WT N2 *C. elegans*, highlighting the functional diversity of proteins across early development and adulthood, as well as in the *aex-3* transgenic background. By combining mass spectrometry with protein–protein interaction (PPI) network analysis and KEGG enrichment, we obtained a systems-level view of how O-GlcNAcylated proteins cluster into functional modules that regulate RNA metabolism, protein folding, and stress responses. These findings suggest that O-GlcNAcylation does not occur randomly across the proteome but preferentially modifies proteins occupying key regulatory nodes.

### O-GlcNAcylation in Developmental Regulation

The protein set in the WT N2 strain was dominated by factors involved in RNA metabolism, ribosomal function, and mechanosensory regulation. In young WT larvae (N2L1). The detection of Q20127, associated with mechanosensory perception and oxidoreductase activity, suggests that even at this early stage, the machinery for sensory input and redox balance is already engaged. Additional proteins such as G5EET4 (glutamic acid-rich protein) and Q4R114 (PIR protein) may act in signalling or structural regulation, though their roles remain less defined. The integrative analysis of the protein–protein interaction (PPI) network and pathway enrichment data revealed that O-GlcNAcylated proteins occupy a non-random, highly interconnected position within the cellular signalling landscape. Rather than being scattered evenly across the proteome, these modified proteins cluster within discrete functional modules, with central hub. Candidate O-GlcNAcylated proteins CWC15 homolog and the ribosomal structural component RL8 underscore the importance of RNA processing and translation during early development. Their clustering in the PPI network supports the notion that O-GlcNAcylation fine-tunes biosynthetic capacity in response to nutrient availability.

In the adult LN2 L4 stage, O-GlcNAc-modified proteins include a KH domain–containing protein, isoform b (Q9BLA0), which is associated with RNA binding (FDR < 0.001) and interactions with miRNA-associated factors, as well as a VWFA domain–containing protein (G5EGU7) involved in protein–protein interactions. Network topology analysis reveals that these proteins have reduced connectivity to ribosomal and spliceosomal modules, indicating a developmental reprogramming of O-GlcNAc signalling. In the adulthood, O-GlcNAcylation shifts from supporting biosynthetic growth to prioritizing the regulation of energy metabolism, proteostasis, and longevity-associated processes in WT *C. elegans* (Tindale et al., 2017; Haskell et al., 2021). This shift aligns with the physiological transition from proliferative capacity to metabolic efficiency and stress resilience. O-GlcNAc modification of FUBP, the human ortholog of Q9BLA0, is linked to the regulation of collagen deposition and the resolution of pulmonary fibrosis (Ding et al., 2017; Vang et al., 2024), indicating that KH domain– containing proteins are likely conserved targets of this modification. InterPro enrichment analysis demonstrates a significant overrepresentation of KH domains, which are conserved RNA-binding modules involved in splicing, stability, transport, and translational regulation. The consistent detection of related InterPro terms across annotation levels reinforces the centrality of RNA-binding via KH domains in the modified protein set. These findings suggest that O-GlcNAcylation modulates the stability, localization, and interactions of KH domain–containing proteins or their RNA substrates, thereby influencing gene expression and cellular responses in the adult worm.

### Crosstalk with Other Post-Translational Modifications

Our proteomic analysis revealed an O-GlcNAcylated peptide with other post-translational modification (PTMs), suggesting complex regulatory interactions that may influence protein function. In particular, the O-GlcNAc-modified peptide **MFITRGLILISLLFVFVMTDDTHDK** annotated for a PIR protein (Q4R114) in the L1N2 strains, was detected with a concurrent methionine oxidation (M1*y2+, neutral loss of m/z 63.99). Such modification is consistent with previously described redox-sensitive PTM sites that influence protein folding or glycosylation efficiency (Khoder-Agha and Kietzmann, 2021). Moreover, in line with findings of Fahie et al. (2022), we proposed that O-GlcNAcylation can intersect with oxidative stress pathways induced by nutrient deprivation, thereby rebuilding or reinforcing the mRNA processing machinery, via spliceosome, to promote cell survival.

### Altered O-GlcNAcylation in the Tauopathy Model

In the *aex-3*p::tau(V337M) strain, O-GlcNAcylation patterns diverged sharply from those in WT N2. At the L1 stage, an extracellular matrix–associated protein such as G5EFA0 was O-GlcNAc modified. This contrasts with the RNA-centric profile observed in WT N2 L1 larvae and suggests that the *aex-3* transgenic strain may differentially regulate extracellular interactions early in development, potentially as a compensatory adaptation to their altered neuronal signalling background.

Conversely, PDI-2, an ER-resident chaperone, was identified as O-GlcNAcylated candidate in the adult aex-3p::tau(V337M) L4, suggesting a stress-adaptive role. Given reports of ER protein reflux into the cytosol under proteotoxic stress (Sicari et al., 2021; Pierre et al., 2024), we propose that PDI-2 could be modified outside the ER, where O-GlcNAcylation may modulate its chaperone function (Figure 2C). Human studies have linked PDI colocalization with TDP-43 aggregates in ALS (Amyotrophic lateral sclerosis) and AD, supporting a conserved role in mitigating stress granule formation (Soares and Martins 2017; Liu et al., 2024).

### Age-Dependent Disruption of O-GlcNAc Cycling in Tauopathy

Our findings indicate that tau expression disrupts the normal, age-related increase in O-GlcNAcylation observed in WT N2 worms, as previously reported in our *in vivo* tracking work of this PTM in *C. elegans* (Garcia, 2025). This impairment may result from competition between phosphorylated tau and O-GlcNAc transferase (OGT) (Huang et al., 2020), which reduces the O-GlcNAc modification of spliceosomal factors and alters mRNA processing. The observed loss of O-GlcNAcylated spliceosomal proteins in adult tau transgenic nematodes suggests inhibition of OGT activity. This inhibition may have downstream effects on intron retention and O-GlcNAc homeostasis (Hsieh, 2019; Cui et al., 2024). These molecular disruptions could intensify proteotoxic stress and contribute to disease progression.

### Broader Implications

Together, these findings support a model in which O-GlcNAcylation acts as a context-dependent regulator of cellular physiology. During development, O-GlcNAcylation coordinates RNA metabolism and translation to promote rapid cellular growth. In adulthood, its role shifts toward maintaining proteostasis and increasing stress resistance. In the context of tauopathy, these regulatory patterns are altered, as indicated by the absence of detectable O-GlcNAc modification of spliceosomal factors. It is proposed that O-GlcNAcylation of pdi-2 occurs in the cytosol in response to redox or/and proteotoxic stress in the ER. This alteration may affect the balance between cellular adaptation and dysfunction.

By mapping these stage- and pathology-specific modifications, our study provides a foundation for exploring O-GlcNAcylation as both a biomarker and a therapeutic target in ageing and neurodegeneration (Huang et al., 2020; Vang et al., 2024)

### Conclusion and Future Directions

Our study reveals that proteomic O-GlcNAcylation patterns in *C. elegans* are dynamically regulated across developmental stages and further reshaped in the context of tau-induced neurodegeneration. WT N2 worms undergo a clear transition from RNA processing and translation in L1 larvae to broader regulation of RNA metabolism, protein stability, and structural remodelling in adults. In contrast, *aex-3*p::tau(V337M) worms exhibit distinct features involving extracellular interactions and stress-related pathways, suggesting compensatory adaptations to altered neuronal signalling and proteotoxic stress.

O-GlcNAcylated PDI-2 protein in the AD model likely regulates protein quality control and ER to cytosol communication under stress. Disrupted age-dependent enhancement of O-GlcNAc levels in tau transgenic worms indicates competition between hyperphosphorylated tau and O-GlcNAc transferase, which provides mechanistic insight into impaired splicing and proteostasis during disease progression. These findings support a model in which O-GlcNAcylation acts as a context-sensitive modulator, adjusting its targets according to developmental stage, cellular stress, and pathological burden.

Future research should focus on experimentally validating specific O-GlcNAc modifications, determining their biochemical effects on RNA metabolism and protein folding, and testing whether pharmacological modulation of O-GlcNAc cycling can restore cellular resilience in ageing and neurodegenerative contexts. Such investigations may facilitate the development of O-GlcNAc–based therapeutic strategies targeting tauopathy and other protein misfolding disorders.

### Limitations

The *aex-3*p::tau(V337M) *C. elegans* strain provides a robust model for tau-driven pathology, but several limitations must be considered. Transgenic lines exhibit reduced lifespan and progeny viability compared to WT N2, complicating population synchronization and long-term experiments (Holcom et al., 2024). Although our proteomic approach achieved broad coverage, candidate peptides with lower ion scores require cautious interpretation and should be validated using targeted analytical methods. Cross-species extrapolation is limited by species-specific differences in O-GlcNAc cycling and neuronal physiology, despite the identification of conserved orthologs. Further investigation is necessary to clarify the causal relationships between O-GlcNAcylation and other post-translational modifications, such as phosphorylation and oxidation.

## Abbreviations

AD: Alzheimer’s disease
Aβ: β-amyloid
BP: Biological process
CC: Cellular component
CWC15_CAEEL: spliceosome associated protein
KEGG: Kyoto Encyclopedia of Genes and Genomes
MF: molecular function
O-GlcNAcylation: O-linked β-N-acetylglucosaminylation
PDI-2: Protein disulfide-isomerase 2
sWGA: succinylated wheat germ agglutinin
VWFA: von Willerbrand factor type A-domain containing protein
WBP: Worm base phenotype

## Acknowledgments

We thank Dr. Thomas Timm for providing technical support in the nano-LC-mass spectrometry method. *aex-3*p::4R1N-Tau(V337M) strain was kindly provided by Dr. Brian C. Kraemer from the UW Division of Gerontology & Geriatric Medicine, Geriatric Research, Education and Clinical Center (GRECC), Department of Medicine, and Division of Neurogenetics, Department of Neurology, University of Washington, Seattle, WA, USA.

## Data and resource availability

Datasets generated and analyzed are provided as Supplementary Data Files 1-3. Additional data are available from the corresponding author upon reasonable request.

## Footnotes

1 Global action plan on the public health response to dementia 2017–2025. Geneva: World Health Organization; 2017. Licence: CC BY-NC-SA 3.0 IGO.

