## Supplementary material for "Proteome-based investigation of O-GlcNAcylation in a *C. elegans* model of Ageing and Alzheimer’s disease: Functional Support for Earlier Hypothesis-Generating Findings": Supplementary_Dataset2.pdf

### Mascot Search Results

#### Peptide View

MS/MS Fragmentation of **PNSRHDNVSPSK**
Found in **G5EET4** in **UP1940_C_elegans**, Glutamic acid-rich protein OS=Caenorhabditis elegans OX=6239 GN=CELE_F40G12.11 PE=1 SV=1

Match to Query 2295: 1742.813688 from(872.414120,2+) intensity(212588.84) scans(40241) rawscans(sn40241) rtinseconds(13366.562) index(1545)
Title: 1546: Scan 40241 (rt=13366.6) [D:\L1N2-01.raw]
Data file L1N2-01.temp.mgf

observedPNy11a1Sy10a2Ry9a3Hy8a4Dy7a5Ny6a6Vy5a7HESy4a8Py3a9HESy2a10Ky1a11y(11)-4061240.5968 | 4.9 ppmy(9)-4061039.5260 | 1.9 ppmy(10)-4061126.5592 | 0.7 ppma(8)892.4515 | -14.7 ppma(7)793.3872 | -21.8 ppma(5)564.3183 | -32.3 ppma(6)679.3577 | -45.1 ppmZoom...600800100012001400m/z020406080100% of base peak0500100015002000250030003500ion current

to

**Monoisotopic mass of neutral peptide Mr(calc):** 1742.8071
**Fixed modifications:** Carbamidomethyl (C) (apply to specified residues or termini only)
**Variable modifications:**
S9 : HexNAc (ST), with neutral losses 203.0794(shown in table), 0.0000
S11 : HexNAc (ST), with neutral losses 203.0794(shown in table), 0.0000
**Ions Score:** 41 **Expect:** 0.00093
**Peak matches:** 7/192 fragment ions using 8 most intense peaks
**Annotated fragments:** 7/192 (help)

| # | a | $a^{++}$ | a* | $a^{*++}$ | b | $b^{++}$ | b* | $b^{*++}$ | Seq. | y | $y^{++}$ | y* | $y^{*++}$ | # |
|---|---|---|---|---|---|---|---|---|---|---|---|---|---|---|
| 1 | 70.0651 | 35.5362 | | | 98.0600 | 49.5337 | | | P | | | | | 12 |
| 2 | 184.1081 | 92.5577 | 167.0815 | 84.0444 | 212.1030 | 106.5551 | 195.0764 | 98.0418 | N | ***1240.6029*** | 620.8051 | 1223.5763 | 612.2918 | 11 |
| 3 | 271.1401 | 136.0737 | 254.1135 | 127.5604 | 299.1350 | 150.0711 | 282.1084 | 141.5579 | S | ***1126.5600*** | 563.7836 | 1109.5334 | 555.2703 | 10 |
| 4 | 427.2412 | 214.1242 | 410.2146 | 205.6110 | 455.2361 | 228.1217 | 438.2096 | 219.6084 | R | ***1039.5279*** | 520.2676 | 1022.5014 | 511.7543 | 9 |
| 5 | ***564.3001*** | 282.6537 | 547.2736 | 274.1404 | 592.2950 | 296.6511 | 575.2685 | 288.1379 | H | 883.4268 | 442.2170 | 866.4003 | 433.7038 | 8 |
| 6 | ***679.3270*** | 340.1672 | 662.3005 | 331.6539 | 707.3220 | 354.1646 | 690.2954 | 345.6513 | D | 746.3679 | 373.6876 | 729.3414 | 365.1743 | 7 |
| 7 | ***793.3700*** | 397.1886 | 776.3434 | 388.6754 | 821.3649 | 411.1861 | 804.3383 | 402.6728 | N | 631.3410 | 316.1741 | 614.3144 | 307.6608 | 6 |
| 8 | ***892.4384*** | 446.7228 | 875.4118 | 438.2096 | 920.4333 | 460.7203 | 903.4068 | 452.2070 | V | 517.2980 | 259.1527 | 500.2715 | 250.6394 | 5 |
| 9 | 979.4704 | 490.2388 | 962.4439 | 481.7256 | 1007.4653 | 504.2363 | 990.4388 | 495.7230 | S | 418.2296 | 209.6184 | 401.2031 | 201.1052 | 4 |
| 10 | 1076.5232 | 538.7652 | 1059.4966 | 530.2520 | 1104.5181 | 552.7627 | 1087.4915 | 544.2494 | P | 331.1976 | 166.1024 | 314.1710 | 157.5892 | 3 |
| 11 | 1163.5552 | 582.2812 | 1146.5287 | 573.7680 | 1191.5501 | 596.2787 | 1174.5236 | 587.7654 | S | 234.1448 | 117.5761 | 217.1183 | 109.0628 | 2 |
| 12 | | | | | | | | | K | 147.1128 | 74.0600 | 130.0863 | 65.5468 | 1 |

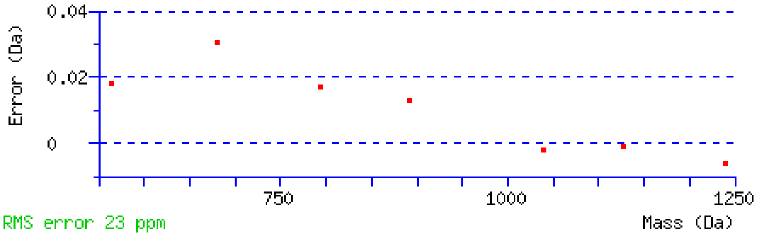


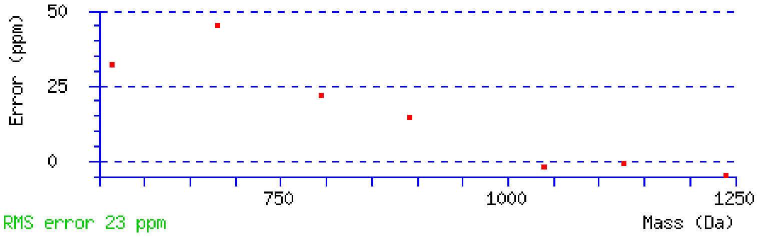


NCBI **BLAST** search of PNSRHDNVSPSK
(Parameters: blastp, nr protein database, expect=20000, no filter, PAM30)
Other BLAST web gateways

##### All matches to this query

| Score | Mr(calc) | Delta | Sequence | Site Analysis |
|---|---|---|---|---|

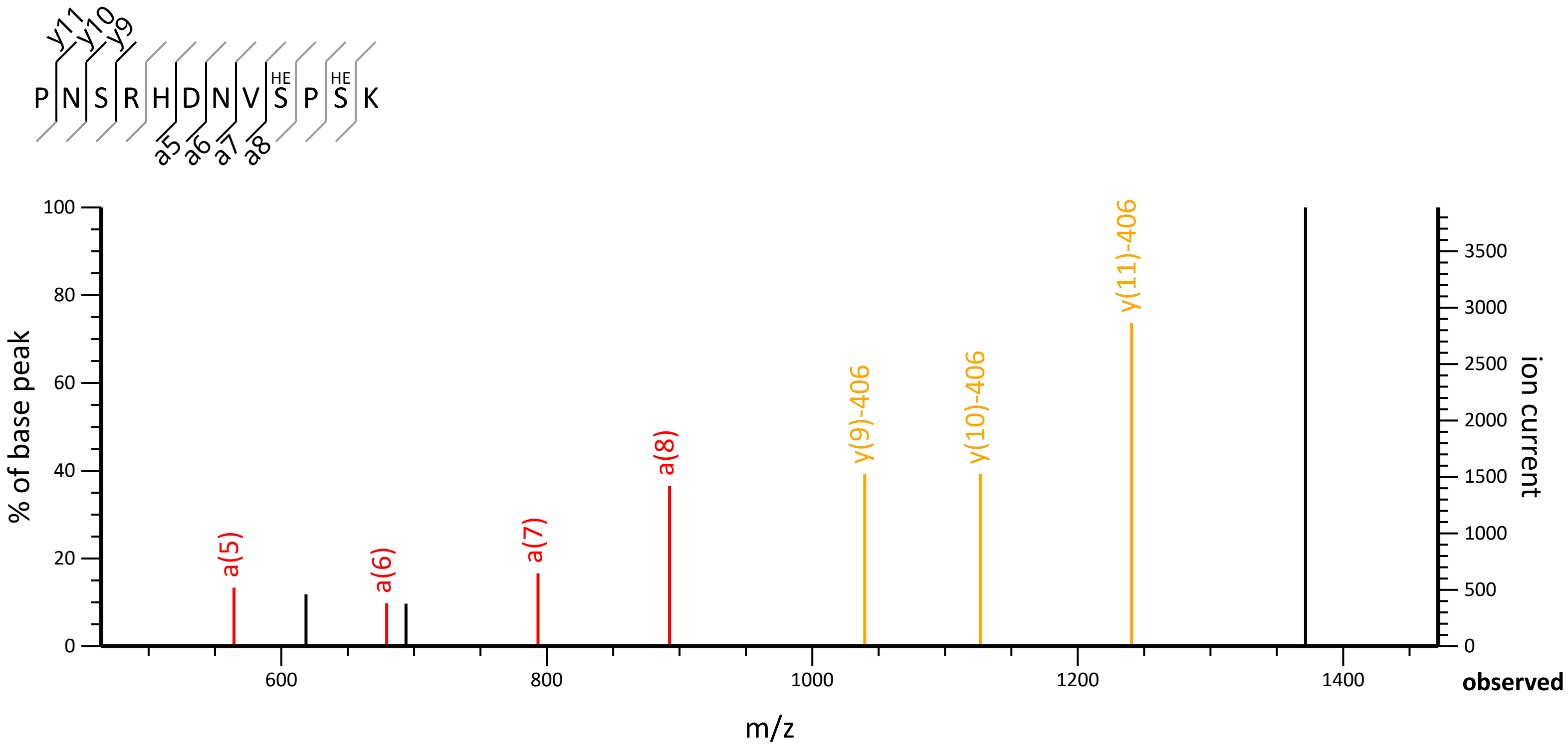

P N S R H D N V S P S K
HE
HE
y11
y10
y9
a5
a6
a7
a8
% of base peak
ion current
m/z
observed
a(5)
a(6)
a(7)
a(8)
y(9)-406
y(10)-406
y(11)-406

| | | | | |
|---|---|---|---|---|
| 40.6 | 1742.8071 | 0.0066 | PNSRHDNVSPSK | HexNAc S9, S11 33.33% |
| 40.6 | 1742.8071 | 0.0066 | PNSRHDNVSPSK | HexNAc S3, S11 33.33% |
| 40.6 | 1742.8071 | 0.0066 | PNSRHDNVSPSK | HexNAc S3, S9 33.33% |
| 7.7 | 1740.8539 | 1.9598 | SKGIPICMVTSGGYQK | |
| 6.9 | 1742.7974 | 0.0163 | NWMSNVAWEFLK | |
| 6.5 | 1740.7836 | 2.0301 | KRTDSEMSQEPSK | |
| 6.5 | 1740.7836 | 2.0301 | KRTDSEMSQEPSK | |
| 5.6 | 1740.8505 | 1.9632 | QMQLTVFYHISR | |
| 3.4 | 1741.8192 | 0.9945 | KDQMSTEEQKDLYK | |
| 3.3 | 1742.8397 | -0.0260 | VYLTSQLSEMTPR | |

### Mascot Search Results

#### Peptide View

MS/MS Fragmentation of **RARDSASSSSSHSK**
Found in **CWC15_CAEEL** in **SwissProt**, Spliceosome-associated protein CWC15 homolog OS=Caenorhabditis elegans OX=6239 GN=cwc-15 PE=1 SV=1

Match to Query 4021: 2479.047912 from(827.356580,3+) intensity(86791.75) scans(14838) rawscans(sn14838) rtinseconds(5599.9938) index(26)
Title: 27: Scan 14838 (rt=5599.99) [D:\L1N2-01.raw]
Data file L1N2-01.temp.mgf

observedRAy13a1Ry12a2Dy11a3Sy10a4Ay9a5Sy8a6HESy7a7HESy6a8HESy5a9HESy4a10Hy3a11HESy2a12Ky1a13a(11)-8121064.4712 | 34.5 ppma(7)716.3278 | 72.7 ppma(10)-609977.4445 | 32.1 ppma(6)629.2895 | 92.6 ppmy(5)-609545.3331 | -119.8 ppmZoom...6008001000120014001600m/z020406080100% of base peak02004006008001000ion current

to

**Monoisotopic mass of neutral peptide Mr(calc):** 2477.0889
**Fixed modifications:** Carbamidomethyl (C) (apply to specified residues or termini only)
**Variable modifications:**
S8 : HexNAc (ST), with neutral losses 203.0794(shown in table), 0.0000
S9 : HexNAc (ST), with neutral losses 203.0794(shown in table), 0.0000
S10 : HexNAc (ST), with neutral losses 203.0794(shown in table), 0.0000
S11 : HexNAc (ST), with neutral losses 203.0794(shown in table), 0.0000
S13 : HexNAc (ST), with neutral losses 203.0794(shown in table), 0.0000
**Ions Score:** 23 **Expect:** 0.018
**Peak matches:** 4/252 fragment ions using 6 most intense peaks
**Annotated fragments:** 5/252 (help)

| # | a | a++ | a* | a*++ | b | b++ | b* | b*++ | Seq. | y | y++ | y* | y*++ | # |
|---|---|---|---|---|---|---|---|---|---|---|---|---|---|---|
| 1 | 129.1135 | 65.0604 | 112.0869 | 56.5471 | 157.1084 | 79.0578 | 140.0818 | 70.5446 | R | | | | | 14 |
| 2 | 200.1506 | 100.5789 | 183.1240 | 92.0657 | 228.1455 | 114.5764 | 211.1190 | 106.0631 | A | 1306.5982 | 653.8027 | 1289.5716 | 645.2895 | 13 |
| 3 | 356.2517 | 178.6295 | 339.2251 | 170.1162 | 384.2466 | 192.6269 | 367.2201 | 184.1137 | R | 1235.5611 | 618.2842 | 1218.5345 | 609.7709 | 12 |
| 4 | 471.2786 | 236.1430 | 454.2521 | 227.6297 | 499.2736 | 250.1404 | 482.2470 | 241.6271 | D | 1079.4600 | 540.2336 | 1062.4334 | 531.7203 | 11 |
| 5 | 558.3107 | 279.6590 | 541.2841 | 271.1457 | 586.3056 | 293.6564 | 569.2790 | 285.1432 | S | 964.4330 | 482.7202 | 947.4065 | 474.2069 | 10 |
| 6 | ***629.3478*** | 315.1775 | 612.3212 | 306.6643 | 657.3427 | 329.1750 | 640.3161 | 320.6617 | A | 877.4010 | 439.2041 | 860.3744 | 430.6909 | 9 |
| 7 | ***716.3798*** | 358.6935 | 699.3533 | 350.1803 | 744.3747 | 372.6910 | 727.3482 | 364.1777 | S | 806.3639 | 403.6856 | 789.3373 | 395.1723 | 8 |
| 8 | 803.4118 | 402.2096 | 786.3853 | 393.6963 | 831.4068 | 416.2070 | 814.3802 | 407.6937 | S | 719.3319 | 360.1696 | 702.3053 | 351.6563 | 7 |
| 9 | 890.4439 | 445.7256 | 873.4173 | 437.2123 | 918.4388 | 459.7230 | 901.4122 | 451.2098 | S | 632.2998 | 316.6536 | 615.2733 | 308.1403 | 6 |
| 10 | ***977.4759*** | 489.2416 | 960.4493 | 480.7283 | 1005.4708 | 503.2390 | 988.4443 | 494.7258 | S | 545.2678 | 273.1375 | 528.2413 | 264.6243 | 5 |
| 11 | ***1064.5079*** | 532.7576 | 1047.4814 | 524.2443 | 1092.5028 | 546.7551 | 1075.4763 | 538.2418 | S | 458.2358 | 229.6215 | 441.2092 | 221.1082 | 4 |
| 12 | 1201.5668 | 601.2871 | 1184.5403 | 592.7738 | 1229.5617 | 615.2845 | 1212.5352 | 606.7712 | H | 371.2037 | 186.1055 | 354.1772 | 177.5922 | 3 |
| 13 | 1288.5989 | 644.8031 | 1271.5723 | 636.2898 | 1316.5938 | 658.8005 | 1299.5672 | 650.2873 | S | 234.1448 | 117.5761 | 217.1183 | 109.0628 | 2 |
| 14 | | | | | | | | | K | 147.1128 | 74.0600 | 130.0863 | 65.5468 | 1 |

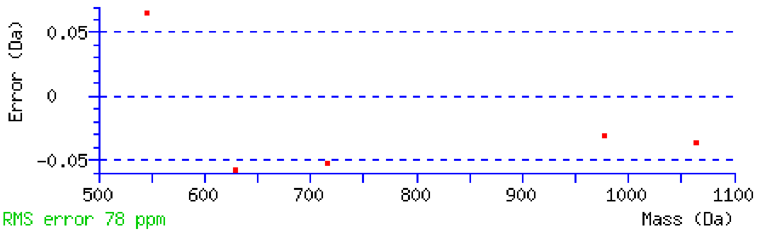


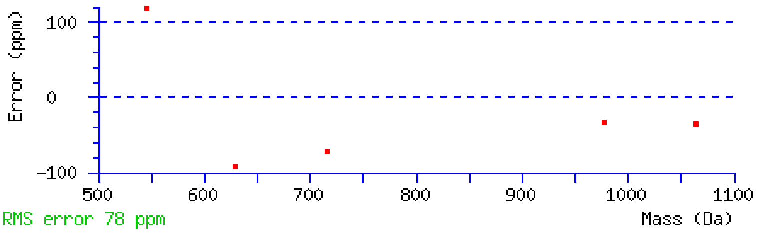


NCBI **BLAST** search of RARDSASSSSSHSK
(Parameters: blastp, nr protein database, expect=20000, no filter, PAM30)
Other BLAST web gateways

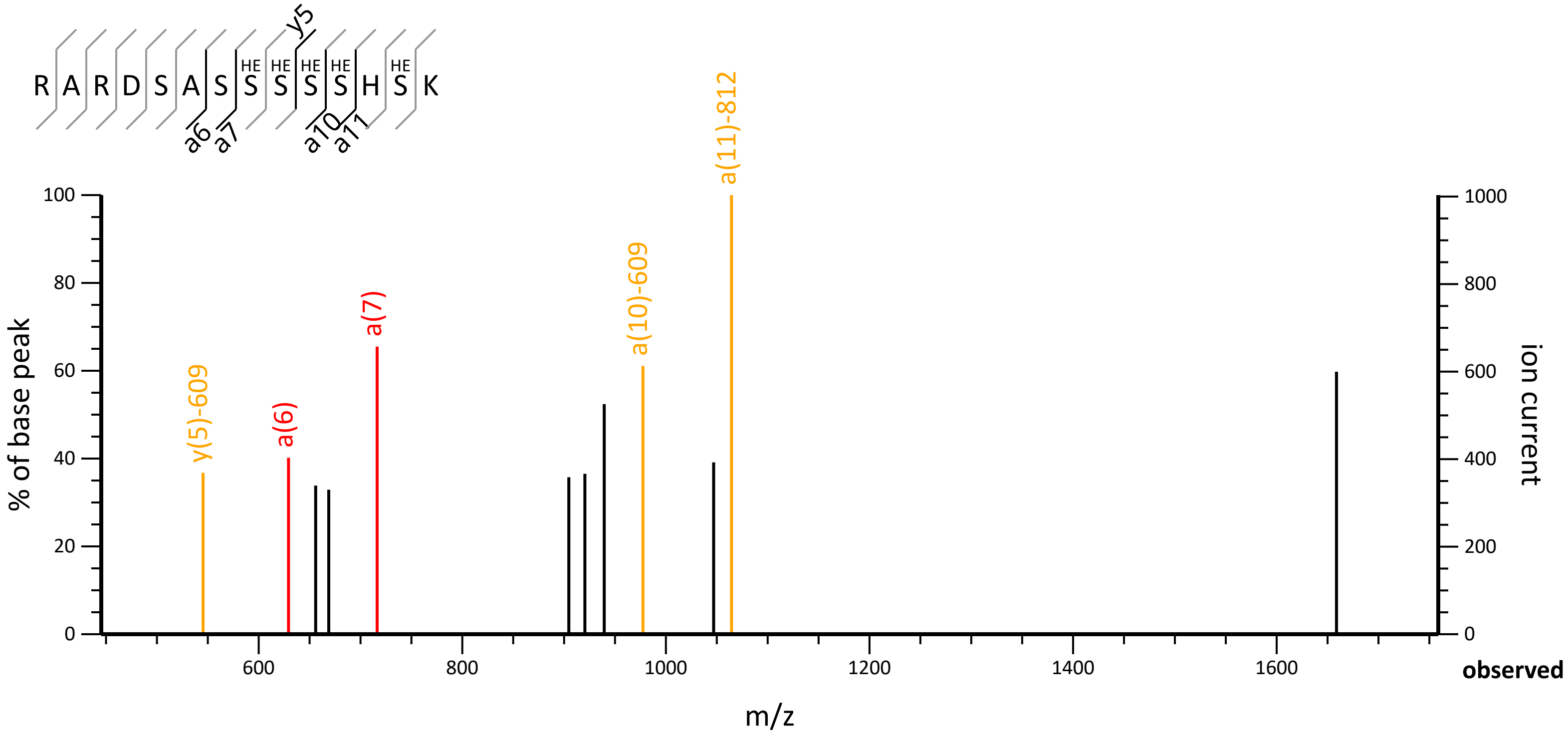
R A R D S A S S S S S H S K
HE HE HE HE HE
y5
a6 a7 a10 a11
y(5)-609
a(6)
a(7)
a(10)-609
a(11)-812
% of base peak
ion current
m/z
observed
0
20
40
60
80
100
200
400
600
800
1000
1200
1400
1600

**All matches to this query**

| Score | Mr(calc) | Delta | Sequence | Site Analysis |
|---|---|---|---|---|
| 22.9 | 2477.0889 | 1.9590 | RARDSASSSSSHSK | HexNAc S8, S9, S10, S11, S13 5.34% |
| 22.9 | 2477.0889 | 1.9590 | RARDSASSSSSHSK | HexNAc S7, S9, S10, S11, S13 5.34% |
| 22.9 | 2477.0889 | 1.9590 | RARDSASSSSSHSK | HexNAc S7, S8, S10, S11, S13 5.34% |
| 22.9 | 2477.0889 | 1.9590 | RARDSASSSSSHSK | HexNAc S7, S8, S9, S11, S13 5.34% |
| 22.9 | 2477.0889 | 1.9590 | RARDSASSSSSHSK | HexNAc S7, S8, S9, S10, S13 5.34% |
| 22.9 | 2477.0889 | 1.9590 | RARDSASSSSSHSK | HexNAc S7, S8, S9, S10, S11 5.34% |
| 22.9 | 2477.0889 | 1.9590 | RARDSASSSSSHSK | HexNAc S5, S9, S10, S11, S13 5.34% |
| 22.9 | 2477.0889 | 1.9590 | RARDSASSSSSHSK | HexNAc S5, S8, S10, S11, S13 5.34% |
| 22.9 | 2477.0889 | 1.9590 | RARDSASSSSSHSK | HexNAc S5, S8, S9, S11, S13 5.34% |
| 22.9 | 2477.0889 | 1.9590 | RARDSASSSSSHSK | HexNAc S5, S8, S9, S10, S13 5.34% |

**Mascot:** http://www.matrixscience.com/

### MATRIX SCIENCE Mascot Search Results

#### Peptide View

MS/MS Fragmentation of **TFDFRADKILESLTNSLK**
Found in **Q20127** in **UP1940_C_elegans**, NADP-dependent oxidoreductase domain-containing protein OS=Caenorhabditis elegans OX=6239 GN=mec-14 PE=4 SV=2

Match to Query 3668: 2301.224472 from(768.082100,3+) intensity(520592.22) scans(40707) rawscans(sn40707) rtinseconds(13456.88) index(1710)
Title: 1711: Scan 40707 (rt=13456.9) [D:\L1N2-01.raw]
Data file L1N2-01.temp.mgf

observedTFy17L1Dy16L2Fy15L3Ry14L4Ay13L5Dy12L6Ky11L7Iy10L8Ly9L9Ey8L10Sy7L11Ly6L12Ty5L13Ny4L14HES y3L15Ly2L16Ky1L17y(7)-203762.4382 | -3.4 ppmy*(6)861.5020 | -52.9 ppmy(10)-2031117.6141 | 28.8 ppmy(5)-203562.3251 | -10.0 ppmy(6)-203675.4118 | -12.1 ppmZoom...500600700800900100011001200m/z020406080100% of base peak05001000150020002500300035000ion current

to

**Monoisotopic mass of neutral peptide Mr(calc):** 2300.1900
**Fixed modifications:** Carbamidomethyl (C) (apply to specified residues or termini only)
**Variable modifications:**
S16 : HexNAc (ST), with neutral losses 203.0794(shown in table), 0.0000
**Ions Score:** 22 **Expect:** 0.046
**Peak matches:** 4/264 fragment ions using 6 most intense peaks
**Annotated fragments:** 5/264 (help)

| # | a | a++ | a* | a*++ | b | b++ | b* | b*++ | Seq. | y | y++ | y* | y*++ | # |
|---|---|---|---|---|---|---|---|---|---|---|---|---|---|---|
| 1 | 74.0600 | 37.5337 | | | 102.0550 | 51.5311 | | | T | | | | | 18 |
| 2 | 221.1285 | 111.0679 | | | 249.1234 | 125.0653 | | | F | 1997.0702 | 999.0387 | 1980.0437 | 990.5255 | 17 |
| 3 | 336.1554 | 168.5813 | | | 364.1503 | 182.5788 | | | D | 1850.0018 | 925.5045 | 1832.9753 | 916.9913 | 16 |
| 4 | 483.2238 | 242.1155 | | | 511.2187 | 256.1130 | | | F | 1734.9749 | 867.9911 | 1717.9483 | 859.4778 | 15 |
| 5 | 639.3249 | 320.1661 | 622.2984 | 311.6528 | 667.3198 | 334.1636 | 650.2933 | 325.6503 | R | 1587.9064 | 794.4569 | 1570.8799 | 785.9436 | 14 |
| 6 | 710.3620 | 355.6847 | 693.3355 | 347.1714 | 738.3570 | 369.6821 | 721.3304 | 361.1688 | A | 1431.8053 | 716.4063 | 1414.7788 | 707.8930 | 13 |
| 7 | 825.3890 | 413.1981 | 808.3624 | 404.6849 | 853.3839 | 427.1956 | 836.3573 | 418.6823 | D | 1360.7682 | 680.8877 | 1343.7417 | 672.3745 | 12 |
| 8 | 953.4839 | 477.2456 | 936.4574 | 468.7323 | 981.4789 | 491.2431 | 964.4523 | 482.7298 | K | 1245.7413 | 623.3743 | 1228.7147 | 614.8610 | 11 |
| 9 | 1066.5680 | 533.7876 | 1049.5415 | 525.2744 | 1094.5629 | 547.7851 | 1077.5364 | 539.2718 | I | ***1117.6463*** | 559.3268 | 1100.6198 | 550.8135 | 10 |
| 10 | 1179.6521 | 590.3297 | 1162.6255 | 581.8164 | 1207.6470 | 604.3271 | 1190.6204 | 595.8139 | L | 1004.5623 | 502.7848 | 987.5357 | 494.2715 | 9 |
| 11 | 1308.6947 | 654.8510 | 1291.6681 | 646.3377 | 1336.6896 | 668.8484 | 1319.6630 | 660.3352 | E | 891.4782 | 446.2427 | 874.4516 | 437.7295 | 8 |
| 12 | 1395.7267 | 698.3670 | 1378.7001 | 689.8537 | 1423.7216 | 712.3644 | 1406.6951 | 703.8512 | S | ***762.4356*** | 381.7214 | 745.4090 | 373.2082 | 7 |
| 13 | 1508.8108 | 754.9090 | 1491.7842 | 746.3957 | 1536.8057 | 768.9065 | 1519.7791 | 760.3932 | L | ***675.4036*** | 338.2054 | 658.3770 | 329.6921 | 6 |
| 14 | 1609.8584 | 805.4329 | 1592.8319 | 796.9196 | 1637.8533 | 819.4303 | 1620.8268 | 810.9170 | T | ***562.3195*** | 281.6634 | 545.2930 | 273.1501 | 5 |
| 15 | 1723.9014 | 862.4543 | 1706.8748 | 853.9410 | 1751.8963 | 876.4518 | 1734.8697 | 867.9385 | N | 461.2718 | 231.1395 | 444.2453 | 222.6263 | 4 |
| 16 | 1810.9334 | 905.9703 | 1793.9068 | 897.4571 | 1838.9283 | 919.9678 | 1821.9018 | 911.4545 | S | 347.2289 | 174.1181 | 330.2023 | 165.6048 | 3 |
| 17 | 1924.0175 | 962.5124 | 1906.9909 | 953.9991 | 1952.0124 | 976.5098 | 1934.9858 | 967.9965 | L | 260.1969 | 130.6021 | 243.1703 | 122.0888 | 2 |
| 18 | | | | | | | | | K | 147.1128 | 74.0600 | 130.0863 | 65.5468 | 1 |

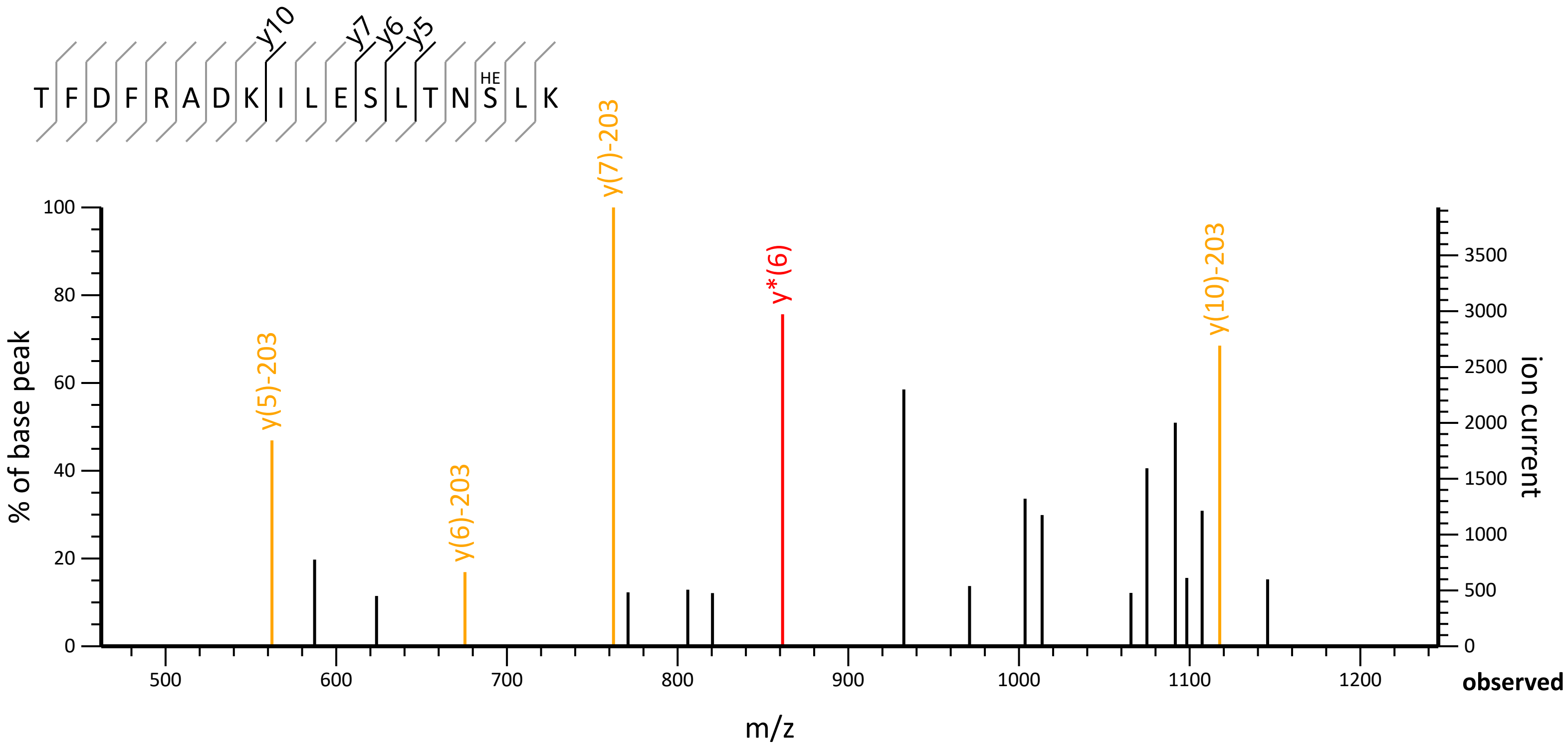
T F D F R A D K I L E S L T N S L K
HE
y10
y7
y6
y5
% of base peak
ion current
y(5)-203
y(6)-203
y(7)-203
y*(6)
y(10)-203
m/z
observed
0
20
40
60
80
100
500
600
700
800
900
1000
1100
1200
0
500
1000
1500
2000
2500
3000
3500

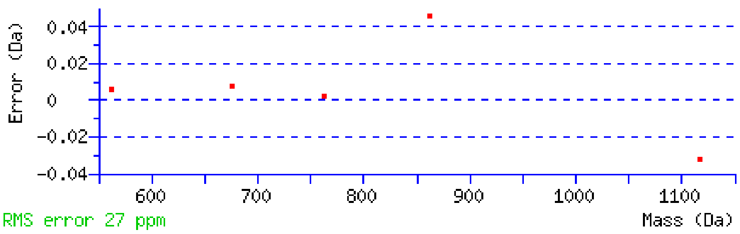


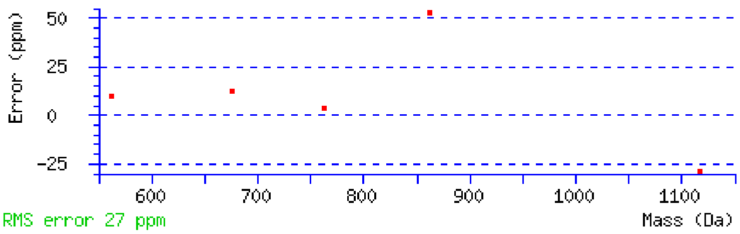


NCBI **BLAST** search of TFDFRADKILESLTNSLK
(Parameters: blastp, nr protein database, expect=20000, no filter, PAM30)
Other BLAST web gateways

**All matches to this query**

| Score | Mr(calc) | Delta | Sequence | Site Analysis |
|---|---|---|---|---|
| 22.2 | 2300.1900 | 1.0345 | TFDFRADKILESLTNSLK | HexNAc S16 25.00% |
| 22.2 | 2300.1900 | 1.0345 | TFDFRADKILESLTNSLK | HexNAc T14 25.00% |
| 22.2 | 2300.1900 | 1.0345 | TFDFRADKILESLTNSLK | HexNAc S12 25.00% |
| 22.2 | 2300.1900 | 1.0345 | TFDFRADKILESLTNSLK | HexNAc T1 25.00% |
| 7.5 | 2299.2172 | 2.0073 | SPILSTIAESIHGASSIRAFDK | |
| 7.2 | 2300.2263 | 0.9982 | LEQLILKNSEELESLQTWK | |
| 7.2 | 2301.2117 | 0.0128 | NAVYNLSLSTLSVNHKIDWK | |
| 5.4 | 2301.1939 | 0.0305 | KRSMSLWVEFITASGYLSAR | |
| 4.7 | 2301.1965 | 0.0280 | ADSILFKTRLPQNHQK | |
| 4.6 | 2299.2199 | 2.0046 | DLFITSLIDTIVKPYR | |

**Mascot:** http://www.matrixscience.com/

### Mascot Search Results

#### Peptide View

MS/MS Fragmentation of **VGLIAARRTGR**
Found in **RL8_CAEEL** in **SwissProt**, Large ribosomal subunit protein uL2 OS=Caenorhabditis elegans OX=6239 GN=rpl-8 PE=1 SV=1

Match to Query 1744: 1373.829788 from(687.922170,2+) intensity(267442.41) scans(42165) rawscans(sn42165) rtinseconds(13713.183) index(2231)
Title: 2232: Scan 42165 (rt=13713.2) [D:\L1N2-01.raw]
Data file L1N2-01.temp.mgf

observedVGy10L1Ly9L2Iy8L3Ay7L4Ay6L5Ry5L6Ry4L7HETy3L8Gy2L9Ry1L10y(7)990.5956 | -52.1 ppmy(6)919.5570 | -54.5 ppmy(5)848.5248 | -65.0 ppmy(8)1103.6871 | -53.6 ppmZoom...600700800900100011001200m/z020406080100% of base peak050010001500ion current

to

```
Monoisotopic mass of neutral peptide Mr(calc): 1371.7946
Fixed modifications: Carbamidomethyl (C) (apply to specified residues or termini only)
Variable modifications:
  T9 : HexNAc (ST), with neutral losses 0.0000(shown in table), 203.0794
Ions Score: 22  Expect: 0.018
Peak matches: 4/144 fragment ions using 8 most intense peaks
Annotated fragments: 4/144   (help)
```

| # | a | a++ | a* | a*++ | b | b++ | b* | b*++ | Seq. | y | y++ | y* | y*++ | # |
|---|---|---|---|---|---|---|---|---|---|---|---|---|---|---|
| 1 | 72.0808 | 36.5440 | | | 100.0757 | 50.5415 | | | V | | | | | 11 |
| 2 | 129.1022 | 65.0548 | | | 157.0972 | 79.0522 | | | G | 1273.7335 | 637.3704 | 1256.7070 | 628.8571 | 10 |
| 3 | 242.1863 | 121.5968 | | | 270.1812 | 135.5942 | | | L | 1216.7120 | 608.8597 | 1199.6855 | 600.3464 | 9 |
| 4 | 355.2704 | 178.1388 | | | 383.2653 | 192.1363 | | | I | ***1103.6280*** | 552.3176 | 1086.6014 | 543.8044 | 8 |
| 5 | 426.3075 | 213.6574 | | | 454.3024 | 227.6548 | | | A | ***990.5439*** | 495.7756 | 973.5174 | 487.2623 | 7 |
| 6 | 497.3446 | 249.1759 | | | 525.3395 | 263.1734 | | | A | ***919.5068*** | 460.2570 | 902.4803 | 451.7438 | 6 |
| 7 | 653.4457 | 327.2265 | 636.4192 | 318.7132 | 681.4406 | 341.2239 | 664.4141 | 332.7107 | R | ***848.4697*** | 424.7385 | 831.4431 | 416.2252 | 5 |
| 8 | 809.5468 | 405.2770 | 792.5203 | 396.7638 | 837.5417 | 419.2745 | 820.5152 | 410.7612 | R | 692.3686 | 346.6879 | 675.3420 | 338.1747 | 4 |
| 9 | 1113.6739 | 557.3406 | 1096.6473 | 548.8273 | 1141.6688 | 571.3380 | 1124.6422 | 562.8248 | T | 536.2675 | 268.6374 | 519.2409 | 260.1241 | 3 |
| 10 | 1170.6953 | 585.8513 | 1153.6688 | 577.3380 | 1198.6902 | 599.8488 | 1181.6637 | 591.3355 | G | 232.1404 | 116.5738 | 215.1139 | 108.0606 | 2 |
| 11 | | | | | | | | | R | 175.1190 | 88.0631 | 158.0924 | 79.5498 | 1 |

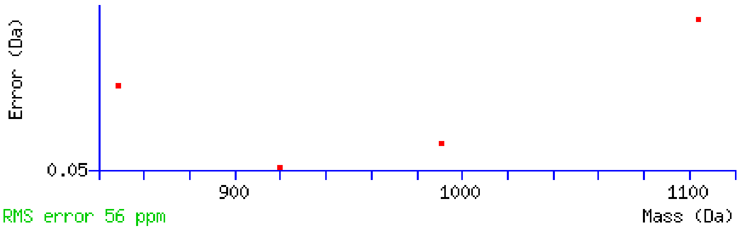


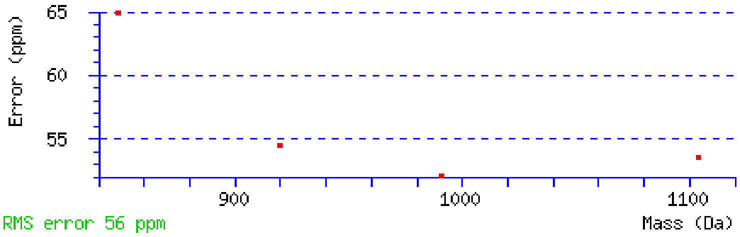


NCBI **BLAST** search of VGLIAARRTGR
(Parameters: blastp, nr protein database, expect=20000, no filter, PAM30)
Other BLAST web gateways

##### All matches to this query

| Score | Mr(calc) | Delta | Sequence |
|---|---|---|---|
| 22.1 | 1371.7946 | 2.0352 | VGLIAARRTGR |
| 3.4 | 1372.8038 | 1.0260 | AKVRQLVTQK |
| 3.0 | 1373.7918 | 0.0379 | LKTLFLAHTK |

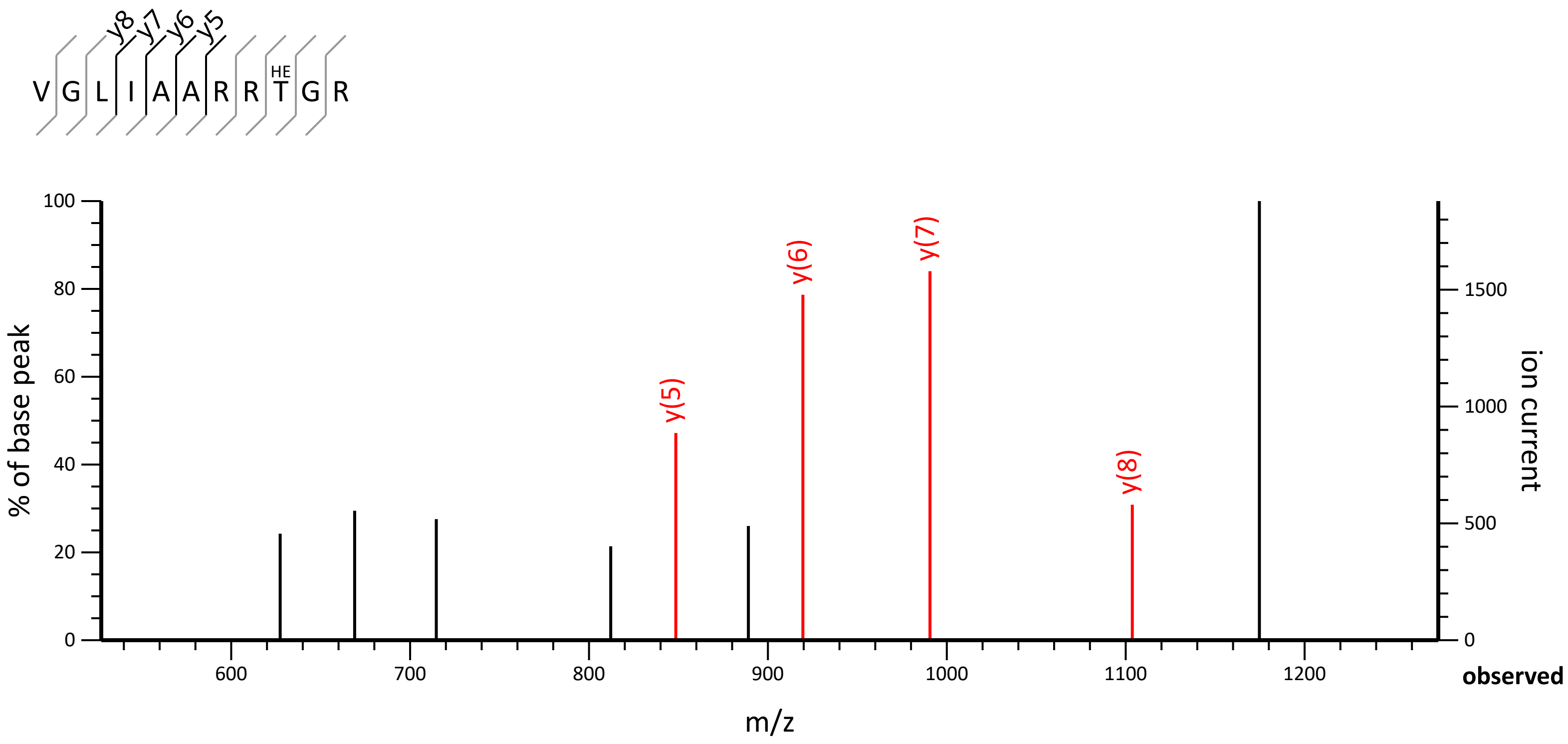

y8 y7 y6 y5
V G L I A A R R T G R
HE
100
80
60
40
20
0
% of base peak
y(5)
y(6)
y(7)
y(8)
1500
1000
500
0
ion current
600
700
800
900
1000
1100
1200
observed
m/z

| | | | |
|---|---|---|---|
| 2.0 | 1371.8020 | 2.0278 | VVAVKTCRIAQK |
| 1.5 | 1373.8395 | -0.0097 | IGTIVNLLPAVHK |
| 0.7 | 1371.8020 | 2.0278 | RVLMLPSVASKR |

**Mascot:** http://www.matrixscience.com/

### MATRIX SCIENCE Mascot Search Results

#### Peptide View

MS/MS Fragmentation of **MFITRGLILISLLFVFVMTDDTHDK**
Found in **Q4R114** in **UP1940_C_elegans**, PIR protein OS=Caenorhabditis elegans OX=6239 GN=CELE_F56D6.12 PE=4 SV=1

Match to Query 5352: 3346.667772 from(1116.563200,3+) intensity(372682.34) scans(41848) rawscans(sn41848) rtinseconds(13659.149) index(2116)
Title: 2117: Scan 41848 (rt=13659.1) [D:\L1N2-01.raw]
Data file L1N2-01.temp.mgf

observedMFy24b1Iy23b2Ty22b3Ry21b4Gy20b5Ly19b6Iy18b7Ly17b8Iy16b9Sy15b10Ly14b11Ly13b12Fy12b13Vy11b14F y10b15Vy9b16OXMy8b17HETy7b18Dy6b19Dy5b20HETy4b21Hy3b22Dy2b23Ky1b24y(15)$^{++}$1095.5348 | -21.2 ppmy(12)$^{++}$938.9871 | -80.5 ppmy(17)$^{++}$1208.6219 | -21.7 ppmb(13)$^{++}$736.3883 | 93.4 ppmy(11)$^{++}$865.4495 | -83.4 ppmZoom...6008001000120014001600m/z020406080100% of base peak010002000300040005000ion current

to

**Monoisotopic mass of neutral peptide Mr(calc):** 3346.7081
**Fixed modifications:** Carbamidomethyl (C) (apply to specified residues or termini only)
**Variable modifications:**
M18 : Oxidation (M), with neutral losses 0.0000(shown in table), 63.9983
T19 : HexNAc (ST), with neutral losses 0.0000(shown in table), 203.0794
T22 : HexNAc (ST), with neutral losses 0.0000(shown in table), 203.0794
**Ions Score:** 22 **Expect:** 0.026
**Peak matches:** 5/644 fragment ions using 6 most intense peaks
**Annotated fragments:** 5/644 (help)

| # | a | a$^{++}$ | a* | a*$^{++}$ | b | b$^{++}$ | b* | b*$^{++}$ | Seq. | y | y$^{++}$ | y* | y*$^{++}$ | # |
|---|---|---|---|---|---|---|---|---|---|---|---|---|---|---|
| 1 | 104.0528 | 52.5301 | | | 132.0478 | 66.5275 | | | M | | | | | 25 |
| 2 | 251.1213 | 126.0643 | | | 279.1162 | 140.0617 | | | F | 3216.6748 | 1608.8411 | 3199.6483 | 1600.3278 | 24 |
| 3 | 364.2053 | 182.6063 | | | 392.2002 | 196.6038 | | | I | 3069.6064 | 1535.3069 | 3052.5799 | 1526.7936 | 23 |
| 4 | 465.2530 | 233.1301 | | | 493.2479 | 247.1276 | | | T | 2956.5224 | 1478.7648 | 2939.4958 | 1470.2515 | 22 |
| 5 | 621.3541 | 311.1807 | 604.3276 | 302.6674 | 649.3490 | 325.1782 | 632.3225 | 316.6649 | R | 2855.4747 | 1428.2410 | 2838.4481 | 1419.7277 | 21 |
| 6 | 678.3756 | 339.6914 | 661.3490 | 331.1782 | 706.3705 | 353.6889 | 689.3439 | 345.1756 | G | 2699.3736 | 1350.1904 | 2682.3470 | 1341.6772 | 20 |
| 7 | 791.4596 | 396.2335 | 774.4331 | 387.7202 | 819.4546 | 410.2309 | 802.4280 | 401.7176 | L | 2642.3521 | 1321.6797 | 2625.3256 | 1313.1664 | 19 |
| 8 | 904.5437 | 452.7755 | 887.5172 | 444.2622 | 932.5386 | 466.7729 | 915.5121 | 458.2597 | I | 2529.2681 | 1265.1377 | 2512.2415 | 1256.6244 | 18 |
| 9 | 1017.6278 | 509.3175 | 1000.6012 | 500.8042 | 1045.6227 | 523.3150 | 1028.5961 | 514.8017 | L | 2416.1840 | ***1208.5956*** | 2399.1574 | 1200.0824 | 17 |
| 10 | 1130.7118 | 565.8596 | 1113.6853 | 557.3463 | 1158.7067 | 579.8570 | 1141.6802 | 571.3437 | I | 2303.0999 | 1152.0536 | 2286.0734 | 1143.5403 | 16 |
| 11 | 1217.7439 | 609.3756 | 1200.7173 | 600.8623 | 1245.7388 | 623.3730 | 1228.7122 | 614.8598 | S | 2190.0159 | ***1095.5116*** | 2172.9893 | 1086.9983 | 15 |
| 12 | 1330.8279 | 665.9176 | 1313.8014 | 657.4043 | 1358.8228 | 679.9151 | 1341.7963 | 671.4018 | L | 2102.9838 | 1051.9956 | 2085.9573 | 1043.4823 | 14 |
| 13 | 1443.9120 | 722.4596 | 1426.8854 | 713.9464 | 1471.9069 | **736.4571** | 1454.8804 | 727.9438 | L | 1989.8998 | 995.4535 | 1972.8732 | 986.9402 | 13 |
| 14 | 1590.9804 | 795.9938 | 1573.9539 | 787.4806 | 1618.9753 | 809.9913 | 1601.9488 | 801.4780 | F | 1876.8157 | ***938.9115*** | 1859.7892 | 930.3982 | 12 |
| 15 | 1690.0488 | 845.5280 | 1673.0223 | 837.0148 | 1718.0437 | 859.5255 | 1701.0172 | 851.0122 | V | 1729.7473 | ***865.3773*** | 1712.7207 | 856.8640 | 11 |
| 16 | 1837.1172 | 919.0623 | 1820.0907 | 910.5490 | 1865.1121 | 933.0597 | 1848.0856 | 924.5464 | F | 1630.6789 | 815.8431 | 1613.6523 | 807.3298 | 10 |
| 17 | 1936.1856 | 968.5965 | 1919.1591 | 960.0832 | 1964.1806 | 982.5939 | 1947.1540 | 974.0806 | V | 1483.6105 | 742.3089 | 1466.5839 | 733.7956 | 9 |
| 18 | 2083.2210 | 1042.1142 | 2066.1945 | 1033.6009 | 2111.2160 | 1056.1116 | 2094.1894 | 1047.5983 | M | 1384.5420 | 692.7747 | 1367.5155 | 684.2614 | 8 |
| 19 | 2387.3481 | 1194.1777 | 2370.3215 | 1185.6644 | 2415.3430 | 1208.1751 | 2398.3165 | 1199.6619 | T | 1237.5066 | 619.2570 | 1220.4801 | 610.7437 | 7 |
| 20 | 2502.3750 | 1251.6912 | 2485.3485 | 1243.1779 | 2530.3700 | 1265.6886 | 2513.3434 | 1257.1753 | D | 933.3796 | 467.1934 | 916.3530 | 458.6802 | 6 |
| 21 | 2617.4020 | 1309.2046 | 2600.3754 | 1300.6914 | 2645.3969 | 1323.2021 | 2628.3703 | 1314.6888 | D | 818.3527 | 409.6800 | 801.3261 | 401.1667 | 5 |
| 22 | 2921.5290 | 1461.2682 | 2904.5025 | 1452.7549 | 2949.5240 | 1475.2656 | 2932.4974 | 1466.7523 | T | 703.3257 | 352.1665 | 686.2992 | 343.6532 | 4 |
| 23 | 3058.5879 | 1529.7976 | 3041.5614 | 1521.2843 | 3086.5829 | 1543.7951 | 3069.5563 | 1535.2818 | H | 399.1987 | 200.1030 | 382.1721 | 191.5897 | 3 |
| 24 | 3173.6149 | 1587.3111 | 3156.5883 | 1578.7978 | 3201.6098 | 1601.3085 | 3184.5833 | 1592.7953 | D | 262.1397 | 131.5735 | 245.1132 | 123.0602 | 2 |
| 25 | | | | | | | | | K | 147.1128 | 74.0600 | 130.0863 | 65.5468 | 1 |

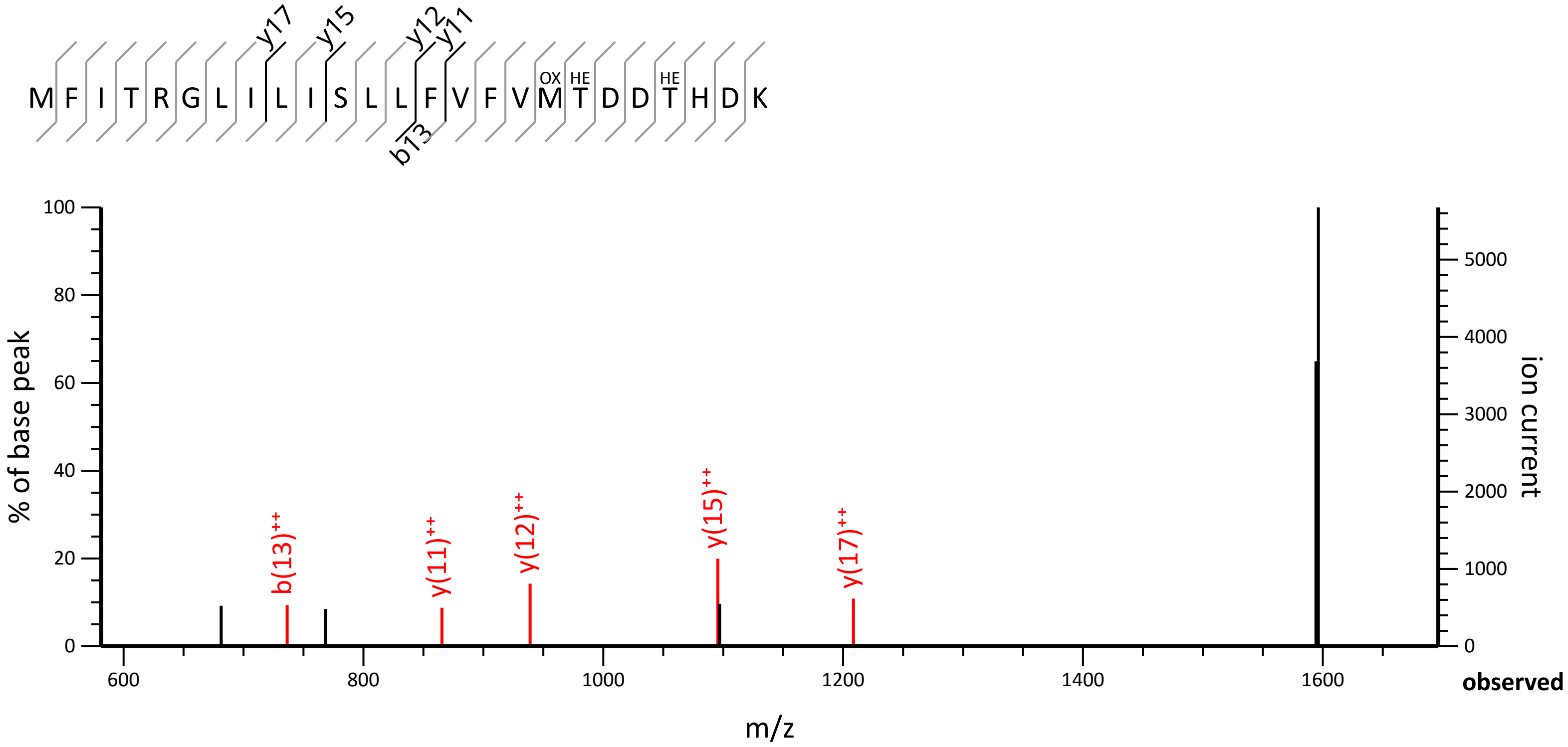
M F I T R G L I L I S L L F V F V M T D D T H D K
OX
HE
HE
y17
y15
y12
y11
b13
% of base peak
ion current
m/z
observed
b(13)++
y(11)++
y(12)++
y(15)++
y(17)++

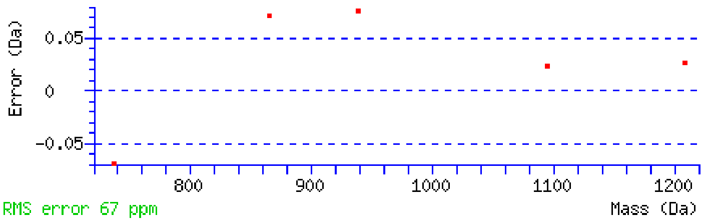


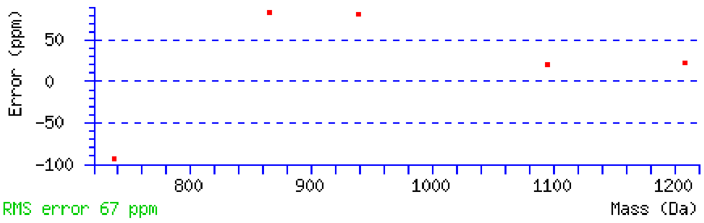


NCBI **BLAST** search of MFITRGLILISLLFVFVMTDDTHDK
(Parameters: blastp, nr protein database, expect=20000, no filter, PAM30)
Other BLAST web gateways

**All matches to this query**

| Score | Mr(calc) | Delta | Sequence | Site Analysis |
|---|---|---|---|---|
| 22.1 | 3346.7081 | -0.0403 | MFITRGLILISLLFVFVMTDDTHDK | HexNAc T19, T22, Oxidation M18; 91.90% |
| 5.0 | 3346.6780 | -0.0102 | RTSHLIKDILDLPTVNGEIDEFGR | |
| 3.2 | 3344.6333 | 2.0344 | NSLISRPPTEKSMAPLGYIDDLYR | |
| 3.2 | 3344.6333 | 2.0344 | NSLISRPPTEKSMAPLGYIDDLYR | |
| 3.2 | 3344.6333 | 2.0344 | NSLISRPPTEKSMAPLGYIDDLYR | |
| 3.2 | 3344.6333 | 2.0344 | NSLISRPPTEKSMAPLGYIDDLYR | |
| 2.7 | 3344.6235 | 2.0443 | HKYTNNENILVDHVEKVDPEVFDIMK | |
| 2.7 | 3346.6941 | -0.0264 | RETLFLTMTVSNGFKAEELMWQSSVSLVK | |
| 2.2 | 3344.6445 | 2.0232 | SADIYSLAVIASEVLTRKEAWNMAER | |
| 2.2 | 3344.6445 | 2.0232 | SADIYSLAVIASEVLTRKEAWNMAER | |

**Mascot:** http://www.matrixscience.com/

### Mascot Search Results

#### Peptide View

MS/MS Fragmentation of **LLVADILACNDDTPASAMMAGNGPVATMSLQVK**
Found in **Q9BLA0** in **UP1940_C_elegans**, K Homology domain-containing protein OS=Caenorhabditis elegans OX=6239 GN=fubl-3 PE=1 SV=1

Match to Query 3660: 3520.028656 from(881.014440,4+) intensity(517471.72) scans(38392) rawscans(sn38392) rtinseconds(13966.695) index(1104)
Title: 1105: Scan 38392 (rt=13966.7) [D:\L4N2-02.raw]
Data file L4N2-02.temp.mgf

Click mouse within plot area to zoom in by factor of two about that point
Or, Plot from 550 to 1300 Da Full range
Show Y-axis

MS/MS spectrum of LLVADILACNDDTPASAMMAGNGPVATMSLQVK

**Monoisotopic mass of neutral peptide Mr(calc):** 3519.6969
**Variable modifications:**
S29 : HexNAc (ST), with neutral losses 0.0000(shown in table), 203.0794
**Ions Score:** 43 **Expect:** 0.011
**Peak matches:** 24/492 fragment ions using 19 most intense peaks
**Annotated fragments:** 26/492 (help)

| # | a | a++ | a* | a*++ | b | b++ | b* | b*++ | Seq. | y | y++ | y* | y*++ | # |
|---|---|---|---|---|---|---|---|---|---|---|---|---|---|---|
| 1 | 86.0964 | 43.5519 | | | 114.0913 | 57.5493 | | | L | | | | | 33 |
| 2 | 199.1805 | 100.0939 | | | 227.1754 | 114.0913 | | | L | 3407.6201 | 1704.3137 | 3390.5936 | 1695.8004 | 32 |
| 3 | 298.2489 | 149.6281 | | | 326.2438 | 163.6255 | | | V | 3294.5361 | 1647.7717 | 3277.5095 | 1639.2584 | 31 |
| 4 | 369.2860 | 185.1466 | | | 397.2809 | 199.1441 | | | A | 3195.4677 | 1598.2375 | 3178.4411 | 1589.7242 | 30 |
| 5 | 484.3130 | 242.6601 | | | 512.3079 | 256.6576 | | | D | 3124.4305 | 1562.7189 | 3107.4040 | 1554.2056 | 29 |
| 6 | 597.3970 | 299.2022 | | | 625.3919 | 313.1996 | | | I | 3009.4036 | 1505.2054 | 2992.3770 | 1496.6922 | 28 |
| 7 | 710.4811 | 355.7442 | | | 738.4760 | 369.7416 | | | L | 2896.3195 | 1448.6634 | 2879.2930 | 1440.1501 | 27 |
| 8 | 781.5182 | 391.2627 | | | 809.5131 | 405.2602 | | | A | 2783.2355 | 1392.1214 | 2766.2089 | 1383.6081 | 26 |
| 9 | **884.5274** | 442.7673 | | | 912.5223 | 456.7648 | | | C | 2712.1984 | 1356.6028 | 2695.1718 | 1348.0895 | 25 |
| 10 | **998.5703** | 499.7888 | 981.5438 | 491.2755 | 1026.5652 | 513.7863 | 1009.5387 | 505.2730 | N | 2609.1892 | 1305.0982 | 2592.1626 | 1296.5849 | 24 |
| 11 | 1113.5973 | 557.3023 | 1096.5707 | 548.7890 | 1141.5922 | 571.2997 | 1124.5656 | 562.7864 | D | 2495.1462 | 1248.0768 | 2478.1197 | 1239.5635 | 23 |
| 12 | 1228.6242 | 614.8157 | 1211.5977 | 606.3025 | 1256.6191 | 628.8132 | 1239.5926 | 620.2999 | D | 2380.1193 | 1190.5633 | 2363.0927 | 1182.0500 | 22 |
| 13 | 1329.6719 | ***665.3396*** | 1312.6453 | 656.8263 | 1357.6668 | 679.3370 | 1340.6402 | 670.8238 | T | 2265.0924 | 1133.0498 | 2248.0658 | 1124.5365 | 21 |
| 14 | 1426.7246 | ***713.8660*** | 1409.6981 | 705.3527 | 1454.7196 | 727.8634 | 1437.6930 | 719.3501 | P | 2164.0447 | 1082.5260 | 2147.0181 | 1074.0127 | 20 |
| 15 | 1497.7618 | 749.3845 | 1480.7352 | 740.8712 | 1525.7567 | ***763.3820*** | 1508.7301 | 754.8687 | A | 2066.9919 | 1033.9996 | 2049.9654 | 1025.4863 | 19 |
| 16 | 1584.7938 | 792.9005 | 1567.7672 | 784.3873 | 1612.7887 | 806.8980 | 1595.7622 | 798.3847 | S | 1995.9548 | **998.4810** | 1978.9282 | 989.9678 | 18 |
| 17 | 1655.8309 | ***828.4191*** | 1638.8044 | 819.9058 | 1683.8258 | 842.4165 | 1666.7993 | 833.9033 | A | 1908.9228 | 954.9650 | 1891.8962 | 946.4517 | 17 |

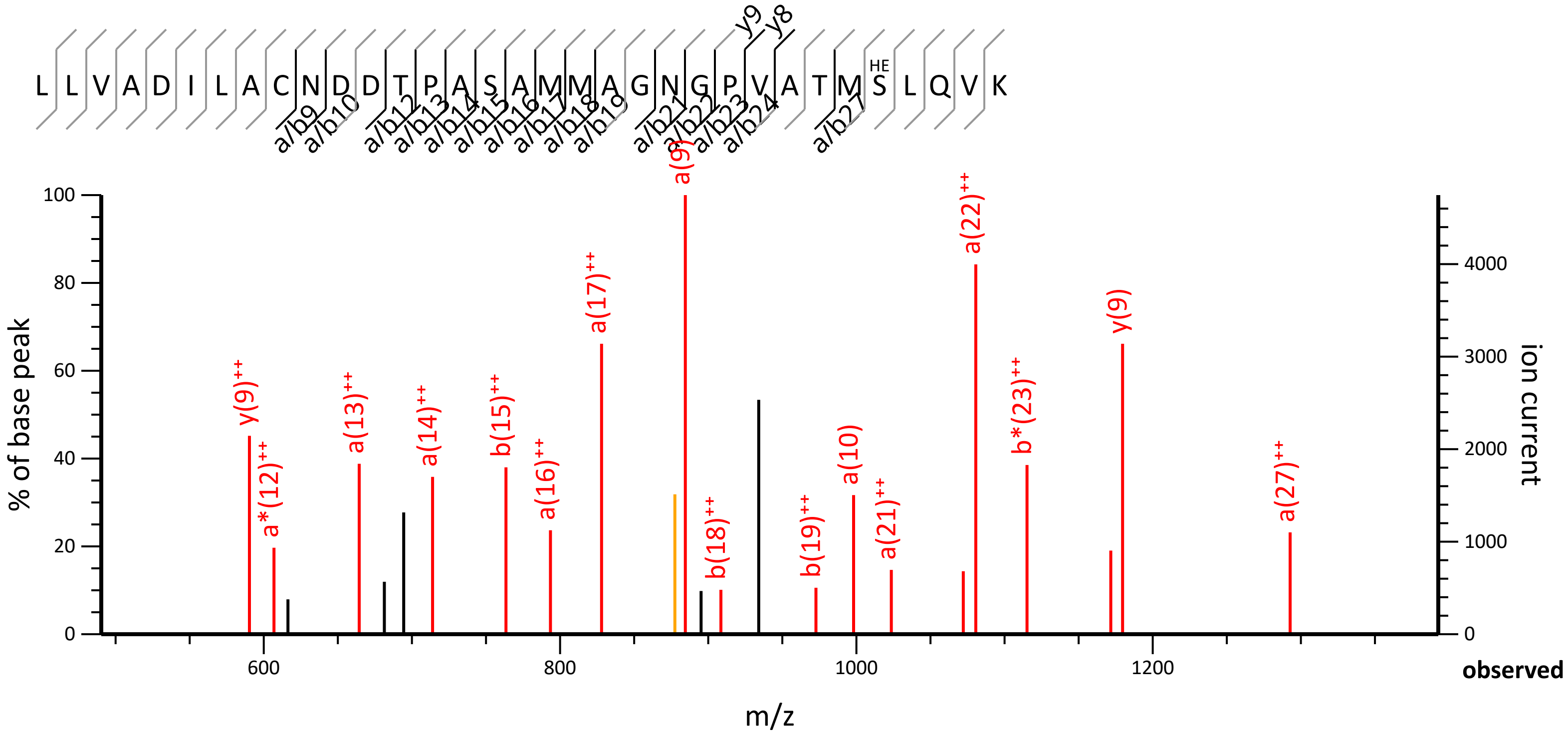

L L V A D I L A C N D D T P A S A M M A G N G P V A T M S L Q V K
HE
y9
y8
a/b9
a/b10
a/b12
a/b13
a/b14
a/b15
a/b16
a/b17
a/b18
a/b19
a/b21
a/b22
a/b23
a/b24
a/b27
% of base peak
ion current
m/z
observed
y(9)++
a*(12)++
a(13)++
a(14)++
b(15)++
a(16)++
a(17)++
a(9)
b(18)++
b(19)++
a(10)
a(21)++
a(22)++
b*(23)++
y(9)
a(27)++
0
20
40
60
80
100
1000
2000
3000
4000
600
800
1000
1200

| | | | | | | | | | | | | | | |
|---|---|---|---|---|---|---|---|---|---|---|---|---|---|---|
| 18 | 1786.8714 | 893.9393 | 1769.8448 | 885.4261 | 1814.8663 | ***907.9368*** | 1797.8397 | 899.4235 | **M** | 1837.8857 | 919.4465 | 1820.8591 | 910.9332 | 16 |
| 19 | 1917.9119 | 959.4596 | 1900.8853 | 950.9463 | 1945.9068 | ***973.4570*** | 1928.8802 | 964.9438 | **M** | 1706.8452 | 853.9262 | 1689.8186 | 845.4129 | 15 |
| 20 | 1988.9490 | 994.9781 | 1971.9224 | 986.4649 | 2016.9439 | 1008.9756 | 1999.9173 | 1000.4623 | **A** | 1575.8047 | 788.4060 | 1558.7781 | 779.8927 | 14 |
| 21 | 2045.9704 | ***1023.4889*** | 2028.9439 | 1014.9756 | 2073.9654 | 1037.4863 | 2056.9388 | 1028.9730 | **G** | 1504.7676 | 752.8874 | 1487.7410 | 744.3741 | 13 |
| 22 | 2160.0134 | ***1080.5103*** | 2142.9868 | 1071.9971 | 2188.0083 | 1094.5078 | 2170.9817 | 1085.9945 | **N** | 1447.7461 | 724.3767 | 1430.7196 | 715.8634 | 12 |
| 23 | 2217.0348 | 1109.0211 | 2200.0083 | 1100.5078 | 2245.0298 | 1123.0185 | 2228.0032 | 1114.5052 | **G** | 1333.7032 | 667.3552 | 1316.6766 | 658.8420 | 11 |
| 24 | 2314.0876 | 1157.5474 | 2297.0611 | 1149.0342 | 2342.0825 | ***1171.5449*** | 2325.0560 | 1163.0316 | **P** | 1276.6817 | 638.8445 | 1259.6552 | 630.3312 | 10 |
| 25 | 2413.1560 | 1207.0816 | 2396.1295 | 1198.5684 | 2441.1509 | 1221.0791 | 2424.1244 | 1212.5658 | **V** | **1179.6290** | **590.3181** | 1162.6024 | 581.8048 | 9 |
| 26 | 2484.1931 | 1242.6002 | 2467.1666 | 1234.0869 | 2512.1880 | 1256.5977 | 2495.1615 | 1248.0844 | **A** | **1080.5605** | 540.7839 | 1063.5340 | 532.2706 | 8 |
| 27 | 2585.2408 | ***1293.1240*** | 2568.2143 | 1284.6108 | 2613.2357 | 1307.1215 | 2596.2092 | 1298.6082 | **T** | 1009.5234 | 505.2654 | 992.4969 | 496.7521 | 7 |
| 28 | 2716.2813 | 1358.6443 | 2699.2547 | 1350.1310 | 2744.2762 | 1372.6417 | 2727.2497 | 1364.1285 | **M** | **908.4757** | 454.7415 | 891.4492 | 446.2282 | 6 |
| 29 | 3006.3927 | 1503.7000 | 2989.3661 | 1495.1867 | 3034.3876 | 1517.6974 | 3017.3611 | 1509.1842 | **S** | 777.4353 | 389.2213 | 760.4087 | 380.7080 | 5 |
| 30 | 3119.4768 | 1560.2420 | 3102.4502 | 1551.7287 | 3147.4717 | 1574.2395 | 3130.4451 | 1565.7262 | **L** | 487.3239 | 244.1656 | 470.2973 | 235.6523 | 4 |
| 31 | 3247.5353 | 1624.2713 | 3230.5088 | 1615.7580 | 3275.5303 | 1638.2688 | 3258.5037 | 1629.7555 | **Q** | 374.2398 | 187.6235 | 357.2132 | 179.1103 | 3 |
| 32 | 3346.6038 | 1673.8055 | 3329.5772 | 1665.2922 | 3374.5987 | 1687.8030 | 3357.5721 | 1679.2897 | **V** | 246.1812 | 123.5942 | 229.1547 | 115.0810 | 2 |
| 33 | | | | | | | | | **K** | 147.1128 | 74.0600 | 130.0863 | 65.5468 | 1 |

Error Distribution

Error Distribution (ppm)

NCBI **BLAST** search of LLVADILACNDDTPASAMMAGNGPVATMSLQVK
(Parameters: blastp, nr protein database, expect=20000, no filter, PAM30)
Other BLAST web gateways

**All matches to this query**

| Score | Mr(calc) | Delta | Sequence | Site Analysis |
|---|---|---|---|---|
| 43.2 | 3519.6969 | 0.3318 | LLVADILACNDDTPASAMMAGNGPVATMSLQVK | HexNAc S29 58.24% |
| 37.2 | 3519.6969 | 0.3318 | LLVADILACNDDTPASAMMAGNGPVATMSLQVK | HexNAc T27 14.33% |
| 37.0 | 3519.6969 | 0.3318 | LLVADILACNDDTPASAMMAGNGPVATMSLQVK | HexNAc S16 13.72% |
| 37.0 | 3519.6969 | 0.3318 | LLVADILACNDDTPASAMMAGNGPVATMSLQVK | HexNAc T13 13.72% |
| 25.0 | 3519.6273 | 0.4014 | GRSFYSEMLLVYCLQSLNSSGPTR | |
| 25.0 | 3519.6273 | 0.4014 | GRSFYSEMLLVYCLQSLNSSGPTR | |
| 24.0 | 3519.7930 | 0.2356 | VLSTEEIEALLKVVETERVAAEAEEAASK | |
| 24.0 | 3519.7930 | 0.2356 | VLSTEEIEALLKVVETERVAAEAEEAASK | |
| 24.0 | 3519.7930 | 0.2356 | VLSTEEIEALLKVVETERVAAEAEEAASK | |
| 24.0 | 3519.7930 | 0.2356 | VLSTEEIEALLKVVETERVAAEAEEAASK | |

**Mascot:** http://www.matrixscience.com/

### Mascot Search Results

#### Peptide View

MS/MS Fragmentation of **HGGTTRTADAIRYATK**
Found in **G5EGU7** in **UP1940_C_elegans**, VWFA domain-containing protein OS=Caenorhabditis elegans OX=6239 GN=C29A12.6 PE=4 SV=1

Match to Query 2435: 2125.062642 from(709.361490,3+) intensity(172035.09) scans(34748) rawscans(sn34748) rtinseconds(13085.666) index(205)
Title: 206: Scan 34748 (rt=13085.7) [D:\L4N2-02.raw]
Data file L4N2-02.temp.mgf

observedHGy15L1Gy14L2Ty13L3Ty12L4Ry11L5HETy10L6Ay9L7Dy8L8Ay7L9Iy6L10Ry5L11Yy4L12Ay3L13HETy2L14K y1L15y(8)-203937.4638 | 49.5 ppmy(9)-2031008.5017 | 45.2 ppmy(6)-203751.4018 | 59.0 ppmy(7)-203822.4363 | 57.0 ppmy(5)-203638.3185 | 68.1 ppmZoom...6007008009001000 1100m/z020406080100% of base peak050010001500200002500ion current

to

**Monoisotopic mass of neutral peptide Mr(calc):** 2124.0447
**Fixed modifications:** Carbamidomethyl (C) (apply to specified residues or termini only)
**Variable modifications:**
T7 : HexNAc (ST), with neutral losses 203.0794(shown in table), 0.0000
T15 : HexNAc (ST), with neutral losses 203.0794(shown in table), 0.0000
**Ions Score:** 34 **Expect:** 0.038
**Peak matches:** 7/288 fragment ions using 4 most intense peaks
**Annotated fragments:** 8/288 (help)

| # | a | a++ | a* | a*++ | b | b++ | b* | b*++ | Seq. | y | y++ | y* | y*++ | # |
|---|---|---|---|---|---|---|---|---|---|---|---|---|---|---|
| 1 | 110.0713 | 55.5393 | | | 138.0662 | 69.5367 | | | H | | | | | 16 |
| 2 | 167.0927 | 84.0500 | | | 195.0877 | 98.0475 | | | G | 1581.8343 | 791.4208 | 1564.8078 | 782.9075 | 15 |
| 3 | 224.1142 | 112.5607 | | | 252.1091 | 126.5582 | | | G | 1524.8129 | 762.9101 | 1507.7863 | 754.3968 | 14 |
| 4 | 325.1619 | 163.0846 | | | 353.1568 | 177.0820 | | | T | 1467.7914 | 734.3993 | 1450.7649 | 725.8861 | 13 |
| 5 | 426.2096 | 213.6084 | | | 454.2045 | 227.6059 | | | T | 1366.7437 | 683.8755 | 1349.7172 | 675.3622 | 12 |
| 6 | 582.3107 | 291.6590 | 565.2841 | 283.1457 | 610.3056 | 305.6564 | 593.2790 | 297.1432 | R | 1265.6961 | 633.3517 | 1248.6695 | 624.8384 | 11 |
| 7 | 683.3583 | 342.1828 | 666.3318 | 333.6695 | 711.3533 | 356.1803 | 694.3267 | 347.6670 | T | 1109.5950 | 555.3011 | 1092.5684 | 546.7878 | 10 |
| 8 | 754.3955 | 377.7014 | 737.3689 | 369.1881 | 782.3904 | 391.6988 | 765.3638 | 383.1856 | A | ***1008.5473*** | 504.7773 | 991.5207 | 496.2640 | 9 |
| 9 | 869.4224 | 435.2148 | 852.3959 | 426.7016 | 897.4173 | 449.2123 | 880.3908 | 440.6990 | D | ***937.5102*** | 469.2587 | 920.4836 | 460.7454 | 8 |
| 10 | 940.4595 | 470.7334 | 923.4330 | 462.2201 | 968.4544 | 484.7309 | 951.4279 | 476.2176 | A | 822.4832 | 411.7452 | 805.4567 | 403.2320 | 7 |
| 11 | 1053.5436 | 527.2754 | 1036.5170 | 518.7622 | 1081.5385 | 541.2729 | 1064.5119 | 532.7596 | I | ***751.4461*** | 376.2267 | 734.4196 | 367.7134 | 6 |
| 12 | 1209.6447 | 605.3260 | 1192.6181 | 596.8127 | 1237.6396 | 619.3234 | 1220.6131 | 610.8102 | R | ***638.3620*** | 319.6847 | 621.3355 | 311.1714 | 5 |
| 13 | 1372.7080 | 686.8577 | 1355.6815 | 678.3444 | 1400.7029 | 700.8551 | 1383.6764 | 692.3418 | Y | 482.2609 | 241.6341 | 465.2344 | 233.1208 | 4 |
| 14 | 1443.7451 | 722.3762 | 1426.7186 | 713.8629 | 1471.7401 | 736.3737 | 1454.7135 | 727.8604 | A | 319.1976 | 160.1024 | 302.1710 | 151.5892 | 3 |
| 15 | 1544.7928 | 772.9000 | 1527.7663 | 764.3868 | 1572.7877 | 786.8975 | 1555.7612 | 778.3842 | T | 248.1605 | 124.5839 | 231.1339 | 116.0706 | 2 |
| 16 | | | | | | | | | K | 147.1128 | 74.0600 | 130.0863 | 65.5468 | 1 |

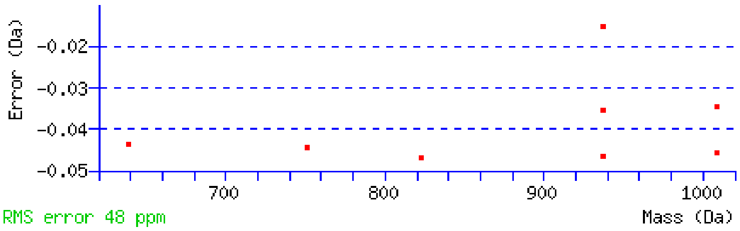


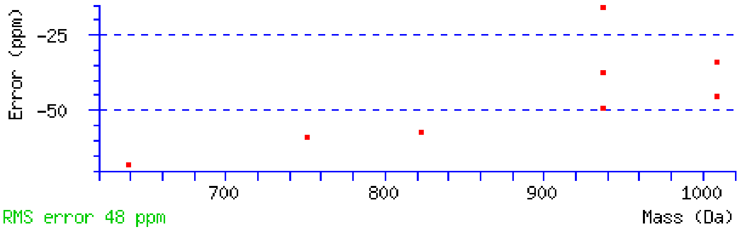


NCBI **BLAST** search of HGGTTRTADAIRYATK

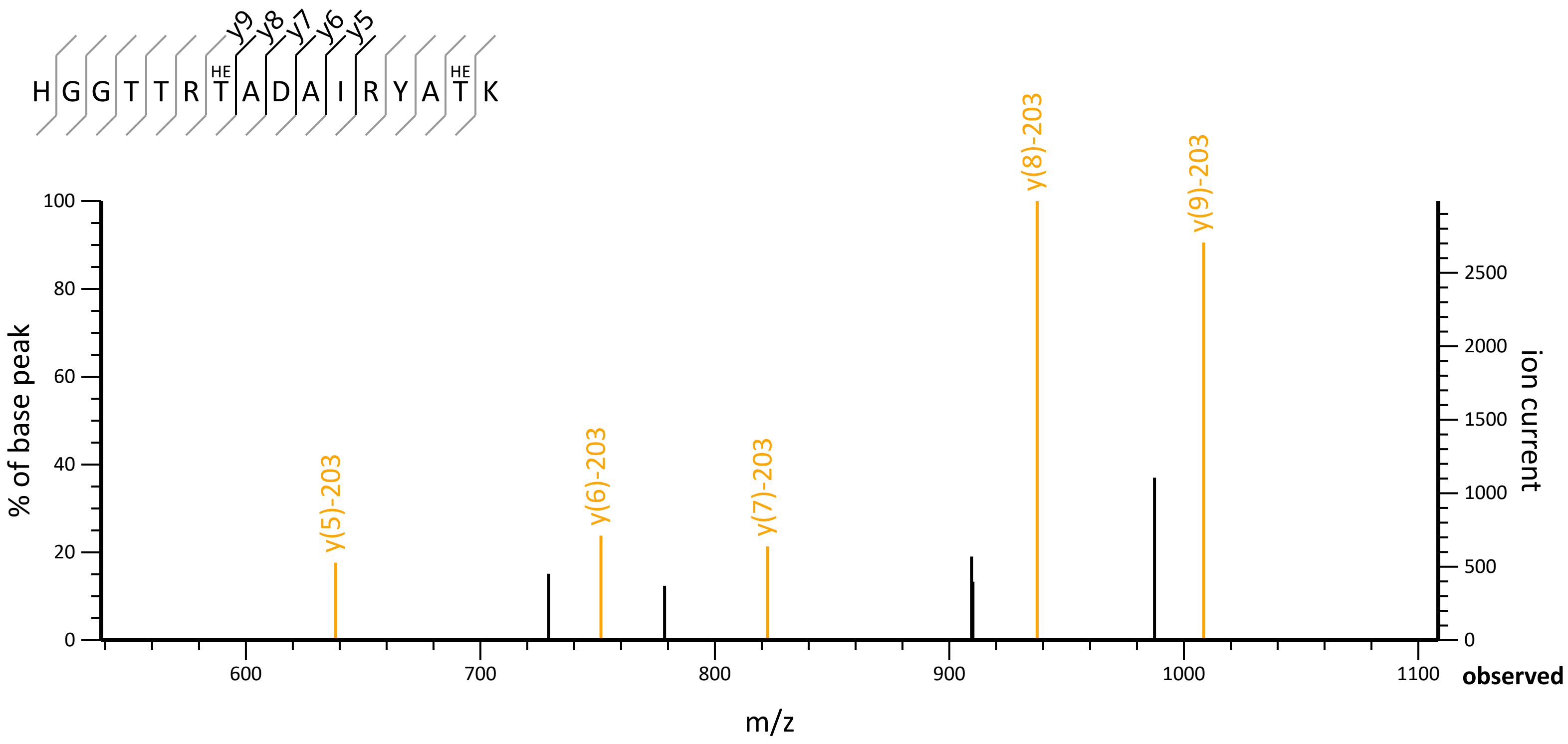

H G G T T R T A D A I R Y A T K
HE
HE
y9 y8 y7 y6 y5
y(5)-203
y(6)-203
y(7)-203
y(8)-203
y(9)-203
% of base peak
ion current
m/z
observed
0
20
40
60
80
100
0
500
1000
1500
2000
2500
600
700
800
900
1000
1100

(Parameters: blastp, nr protein database, expect=20000, no filter, PAM30)
Other BLAST web gateways

**All matches to this query**

| Score | Mr(calc) | Delta | Sequence | Site Analysis |
|---|---|---|---|---|
| 33.6 | 2124.0447 | 1.0179 | HGGTTRTADAIRYATK | HexNAc T7, T15 16.67% |
| 33.6 | 2124.0447 | 1.0179 | HGGTTRTADAIRYATK | HexNAc T5, T15 16.67% |
| 33.6 | 2124.0447 | 1.0179 | HGGTTRTADAIRYATK | HexNAc T5, T7 16.67% |
| 33.6 | 2124.0447 | 1.0179 | HGGTTRTADAIRYATK | HexNAc T4, T15 16.67% |
| 33.6 | 2124.0447 | 1.0179 | HGGTTRTADAIRYATK | HexNAc T4, T7 16.67% |
| 33.6 | 2124.0447 | 1.0179 | HGGTTRTADAIRYATK | HexNAc T4, T5 16.67% |
| 17.3 | 2125.0997 | -0.0370 | FGRYAALSLGVVYGFFR | |
| 16.2 | 2125.0296 | 0.0330 | GGTITTYKDAHNMRVMK | |
| 16.2 | 2125.0296 | 0.0330 | GGTITTYKDAHNMRVMK | |
| 16.2 | 2125.0296 | 0.0330 | GGTITTYKDAHNMRVMK | |

### Mascot Search Results

#### Peptide View

MS/MS Fragmentation of **HGGTTRTADAIRYATK**
Found in **G5EFA0** in **UP1940_C_elegans**, VWFA domain-containing protein OS=Caenorhabditis elegans OX=6239 GN=C29A12.6 PE=4 SV=1

Match to Query 2193: 2125.061592 from(709.361140,3+) intensity(112150.69) scans(34471) rawscans(sn34471) rtinseconds(13079.305) index(124)
Title: 125: Scan 34471 (rt=13079.3) [D:\L1aex3-03.raw]
Data file L1aex3-03.temp.mgf

observedHGy15a/b1Gy14a/b2Ty13a/b3HETy12a/b4Ry11a/b5HETy10a/b6Ay9a/b7Dy8a/b8Ay7a/b9Iy6a/b10Ry5a/b11Yy4a/b12A y3a/b13Ty2a/b14Ky1a/b15y(8)937.4657 | 47.4 ppmy(9)1008.5009 | 46.0 ppma(9)$^{++}$638.3082 | -22.0 ppmy(6)751.4005 | 60.7 ppmb(12)$^{++}$822.4291 | -32.0 ppmy*(12)$^{++}$878.4852 | -49.6 ppmZoom...600800100012001400m/z020406080100% of base peak050010001500ion current

to

**Monoisotopic mass of neutral peptide Mr(calc):** 2124.0447
**Fixed modifications:** Carbamidomethyl (C) (apply to specified residues or termini only)
**Variable modifications:**
T5 : HexNAc (ST), with neutral losses 0.0000(shown in table), 203.0794
T7 : HexNAc (ST), with neutral losses 0.0000(shown in table), 203.0794
**Ions Score:** 30 **Expect:** 0.013
**Peak matches:** 9/268 fragment ions using 10 most intense peaks
**Annotated fragments:** 9/268 (help)

| # | a | a++ | a* | a*++ | b | b++ | b* | b*++ | Seq. | y | y++ | y* | y*++ | # |
|---|---|---|---|---|---|---|---|---|---|---|---|---|---|---|
| 1 | 110.0713 | 55.5393 | | | 138.0662 | 69.5367 | | | H | | | | | 16 |
| 2 | 167.0927 | 84.0500 | | | 195.0877 | 98.0475 | | | G | 1987.9931 | 994.5002 | 1970.9665 | 985.9869 | 15 |
| 3 | 224.1142 | 112.5607 | | | 252.1091 | 126.5582 | | | G | 1930.9716 | 965.9895 | 1913.9451 | 957.4762 | 14 |
| 4 | 325.1619 | 163.0846 | | | 353.1568 | 177.0820 | | | T | 1873.9502 | **937.4787** | 1856.9236 | 928.9654 | 13 |
| 5 | 629.2889 | 315.1481 | | | 657.2838 | 329.1456 | | | T | 1772.9025 | 886.9549 | 1755.8759 | 878.4416 | 12 |
| 6 | 785.3900 | 393.1987 | 768.3635 | 384.6854 | 813.3850 | 407.1961 | 796.3584 | 398.6828 | R | 1468.7754 | 734.8914 | 1451.7489 | 726.3781 | 11 |
| 7 | 1089.5171 | 545.2622 | 1072.4905 | 536.7489 | 1117.5120 | 559.2596 | 1100.4855 | 550.7464 | T | 1312.6743 | 656.8408 | 1295.6478 | 648.3275 | 10 |
| 8 | 1160.5542 | 580.7807 | 1143.5277 | 572.2675 | 1188.5491 | 594.7782 | 1171.5226 | 586.2649 | A | ***1008.5473*** | 504.7773 | 991.5207 | 496.2640 | 9 |
| 9 | 1275.5812 | **638.2942** | 1258.5546 | 629.7809 | 1303.5761 | 652.2917 | 1286.5495 | 643.7784 | D | ***937.5102*** | 469.2587 | 920.4836 | 460.7454 | 8 |
| 10 | 1346.6183 | 673.8128 | 1329.5917 | 665.2995 | 1374.6132 | 687.8102 | 1357.5866 | 679.2970 | A | ***822.4832*** | 411.7452 | 805.4567 | 403.2320 | 7 |
| 11 | 1459.7023 | 730.3548 | 1442.6758 | 721.8415 | 1487.6972 | 744.3523 | 1470.6707 | 735.8390 | I | ***751.4461*** | 376.2267 | 734.4196 | 367.7134 | 6 |
| 12 | 1615.8034 | 808.4054 | 1598.7769 | 799.8921 | 1643.7984 | **822.4028** | 1626.7718 | 813.8895 | R | ***638.3620*** | 319.6847 | 621.3355 | 311.1714 | 5 |
| 13 | 1778.8668 | 889.9370 | 1761.8402 | 881.4237 | 1806.8617 | 903.9345 | 1789.8351 | 895.4212 | Y | 482.2609 | 241.6341 | 465.2344 | 233.1208 | 4 |
| 14 | 1849.9039 | 925.4556 | 1832.8773 | 916.9423 | 1877.8988 | 939.4530 | 1860.8722 | 930.9398 | A | 319.1976 | 160.1024 | 302.1710 | 151.5892 | 3 |
| 15 | 1950.9516 | 975.9794 | 1933.9250 | 967.4661 | 1978.9465 | 989.9769 | 1961.9199 | 981.4636 | T | 248.1605 | 124.5839 | 231.1339 | 116.0706 | 2 |
| 16 | | | | | | | | | K | 147.1128 | 74.0600 | 130.0863 | 65.5468 | 1 |

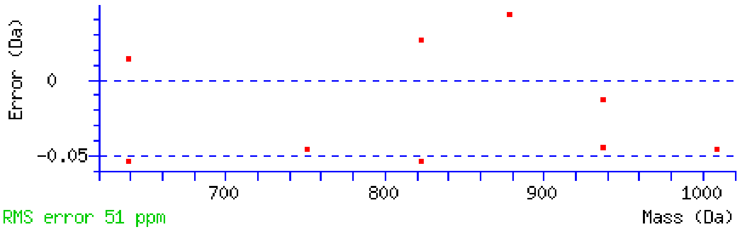


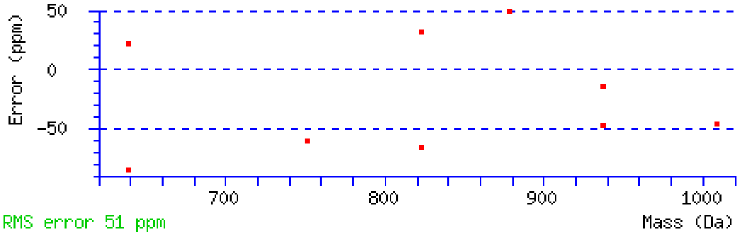


NCBI **BLAST** search of HGGTTRTADAIRYATK

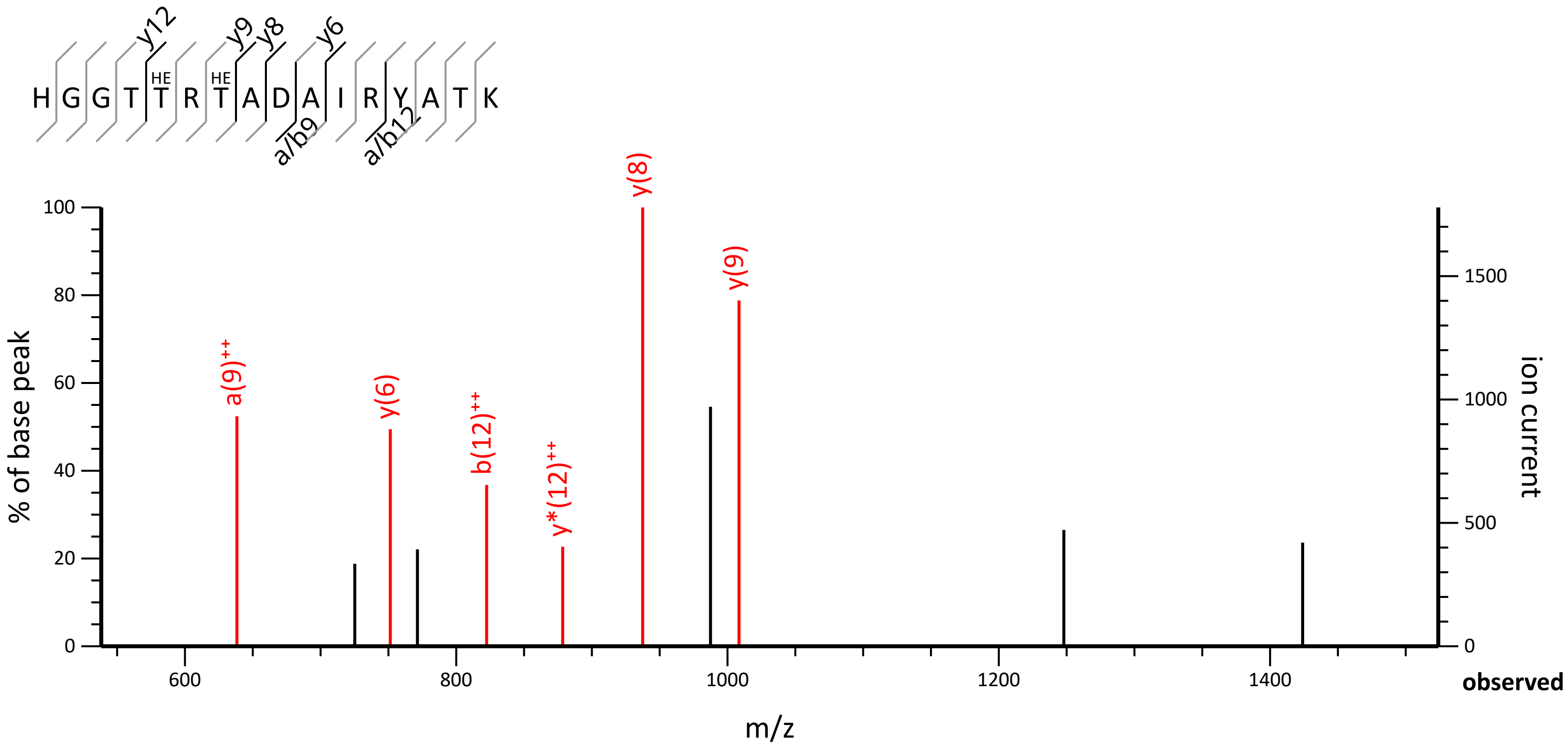

H G G T T R T A D A I R Y A T K
HE
HE
y12
y9
y8
y6
a/b9
a/b12
a(9)++
y(6)
b(12)++
y*(12)++
y(8)
y(9)
% of base peak
ion current
m/z
observed
100
80
60
40
20
0
1500
1000
500
600
800
1000
1200
1400

(Parameters: blastp, nr protein database, expect=20000, no filter, PAM30)
Other BLAST web gateways

**All matches to this query**

| Score | Mr(calc) | Delta | Sequence | Site Analysis |
|---|---|---|---|---|
| 29.8 | 2124.0447 | 1.0169 | HGGTTRTADAIRYATK | HexNAc T5, T7 28.90% |
| 26.7 | 2124.0447 | 1.0169 | HGGTTRTADAIRYATK | HexNAc T7, T15 14.22% |
| 26.7 | 2124.0447 | 1.0169 | HGGTTRTADAIRYATK | HexNAc T5, T15 14.22% |
| 26.7 | 2124.0447 | 1.0169 | HGGTTRTADAIRYATK | HexNAc T4, T15 14.22% |
| 26.7 | 2124.0447 | 1.0169 | HGGTTRTADAIRYATK | HexNAc T4, T7 14.22% |
| 26.7 | 2124.0447 | 1.0169 | HGGTTRTADAIRYATK | HexNAc T4, T5 14.22% |
| 9.0 | 2125.0997 | -0.0381 | FGRYAALSLGVVYGFFR | |
| 7.1 | 2124.0362 | 1.0254 | VVSSAVSTLFNTYK | |
| 7.1 | 2124.0362 | 1.0254 | VVSSAVSTLFNTYK | |
| 6.4 | 2125.0296 | 0.0320 | GGTITTYKDAHNMRVMK | |

**Mascot:** http://www.matrixscience.com/

### Mascot Search Results

#### Peptide View

MS/MS Fragmentation of **DELPAIRLISLEEDMTK**
Found in **PDI2_CAEEL** in **SwissProt**, Protein disulfide-isomerase 2 OS=Caenorhabditis elegans OX=6239 GN=pdi-2 PE=1 SV=1

Match to Query 7478: 2380.180162 from(1191.097357,2+) rtinseconds(14252.2569947) index(6115)
Title: L4aex3-04.39981.39981.2.0.dta
Data file L4aex3-04_HCDFT.mgf

observedDER16a1LR15a2PR14a3AR13a4IR12a5RR11a6LR10a7IR9a8HESR8a9LR7a10ER6a11ER5a12DR4a13MR3a14HET R2a15KR1a16a(13)-203++726.4827 | -101.7 ppma(14)-203++783.9526 | -38.7 ppma(7)++384.3037 | -207.0 ppma(4)++214.1344 | -99.9 ppmZoom...50010001500m/z020406080100% of base peak050010001500200025003000 3500ion current

to

**Monoisotopic mass of neutral peptide Mr(calc):** 2378.1774
**Fixed modifications:** Carboxymethyl (C) (apply to specified residues or termini only)
**Variable modifications:**
S10 : HexNAc (ST), with neutral losses 203.0794(shown in table), 0.0000
T16 : HexNAc (ST), with neutral losses 203.0794(shown in table), 0.0000
**Ions Score:** 22 **Expect:** 0.015
**Peak matches:** 4/284 fragment ions using 6 most intense peaks
**Annotated fragments:** 4/284 (help)

| # | a | a++ | a* | a*++ | b | b++ | b* | b*++ | Seq. | y | y++ | y* | y*++ | # |
|---|---|---|---|---|---|---|---|---|---|---|---|---|---|---|
| 1 | 88.0393 | 44.5233 | | | 116.0342 | 58.5207 | | | D | | | | | 17 |
| 2 | 217.0819 | 109.0446 | | | 245.0768 | 123.0420 | | | E | 1857.9990 | 929.5032 | 1840.9725 | 920.9899 | 16 |
| 3 | 330.1660 | 165.5866 | | | 358.1609 | 179.5841 | | | L | 1728.9564 | 864.9819 | 1711.9299 | 856.4686 | 15 |
| 4 | 427.2187 | ***214.1130*** | | | 455.2136 | 228.1105 | | | P | 1615.8724 | 808.4398 | 1598.8458 | 799.9265 | 14 |
| 5 | 498.2558 | 249.6316 | | | 526.2508 | 263.6290 | | | A | 1518.8196 | 759.9134 | 1501.7931 | 751.4002 | 13 |
| 6 | 611.3399 | 306.1736 | | | 639.3348 | 320.1710 | | | I | 1447.7825 | 724.3949 | 1430.7559 | 715.8816 | 12 |
| 7 | 767.4410 | ***384.2241*** | 750.4145 | 375.7109 | 795.4359 | 398.2216 | 778.4094 | 389.7083 | R | 1334.6984 | 667.8529 | 1317.6719 | 659.3396 | 11 |
| 8 | 880.5251 | 440.7662 | 863.4985 | 432.2529 | 908.5200 | 454.7636 | 891.4934 | 446.2504 | L | 1178.5973 | 589.8023 | 1161.5708 | 581.2890 | 10 |
| 9 | 993.6091 | 497.3082 | 976.5826 | 488.7949 | 1021.6041 | 511.3057 | 1004.5775 | 502.7924 | I | 1065.5133 | 533.2603 | 1048.4867 | 524.7470 | 9 |
| 10 | 1080.6412 | 540.8242 | 1063.6146 | 532.3109 | 1108.6361 | 554.8217 | 1091.6095 | 546.3084 | S | 952.4292 | 476.7182 | 935.4026 | 468.2050 | 8 |
| 11 | 1193.7252 | 597.3663 | 1176.6987 | 588.8530 | 1221.7201 | 611.3637 | 1204.6936 | 602.8504 | L | 865.3972 | 433.2022 | 848.3706 | 424.6889 | 7 |
| 12 | 1322.7678 | 661.8876 | 1305.7413 | 653.3743 | 1350.7627 | 675.8850 | 1333.7362 | 667.3717 | E | 752.3131 | 376.6602 | 735.2865 | 368.1469 | 6 |
| 13 | 1451.8104 | ***726.4088*** | 1434.7839 | 717.8956 | 1479.8053 | 740.4063 | 1462.7788 | 731.8930 | E | 623.2705 | 312.1389 | 606.2440 | 303.6256 | 5 |
| 14 | 1566.8374 | ***783.9223*** | 1549.8108 | 775.4090 | 1594.8323 | 797.9198 | 1577.8057 | 789.4065 | D | 494.2279 | 247.6176 | 477.2014 | 239.1043 | 4 |
| 15 | 1697.8778 | 849.4426 | 1680.8513 | 840.9293 | 1725.8728 | 863.4400 | 1708.8462 | 854.9267 | M | 379.2010 | 190.1041 | 362.1744 | 181.5908 | 3 |
| 16 | 1798.9255 | 899.9664 | 1781.8990 | 891.4531 | 1826.9204 | 913.9639 | 1809.8939 | 905.4506 | T | 248.1605 | 124.5839 | 231.1339 | 116.0706 | 2 |
| 17 | | | | | | | | | K | 147.1128 | 74.0600 | 130.0863 | 65.5468 | 1 |

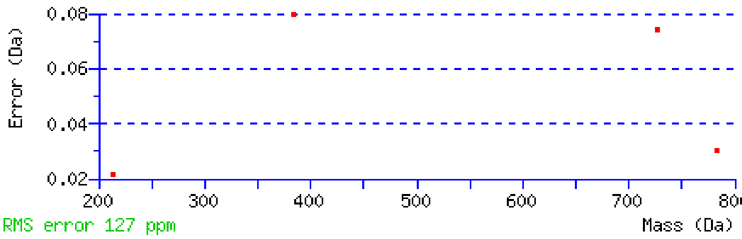


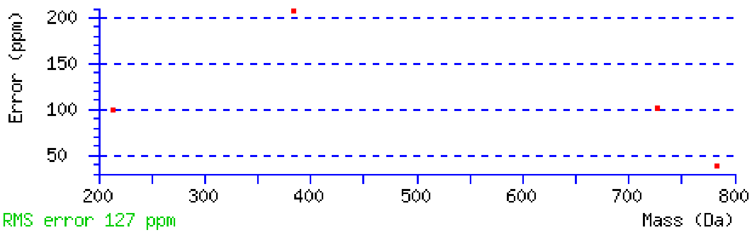


NCBI **BLAST** search of DELPAIRLISLEEDMTK
(Parameters: blastp, nr protein database, expect=20000, no filter, PAM30)

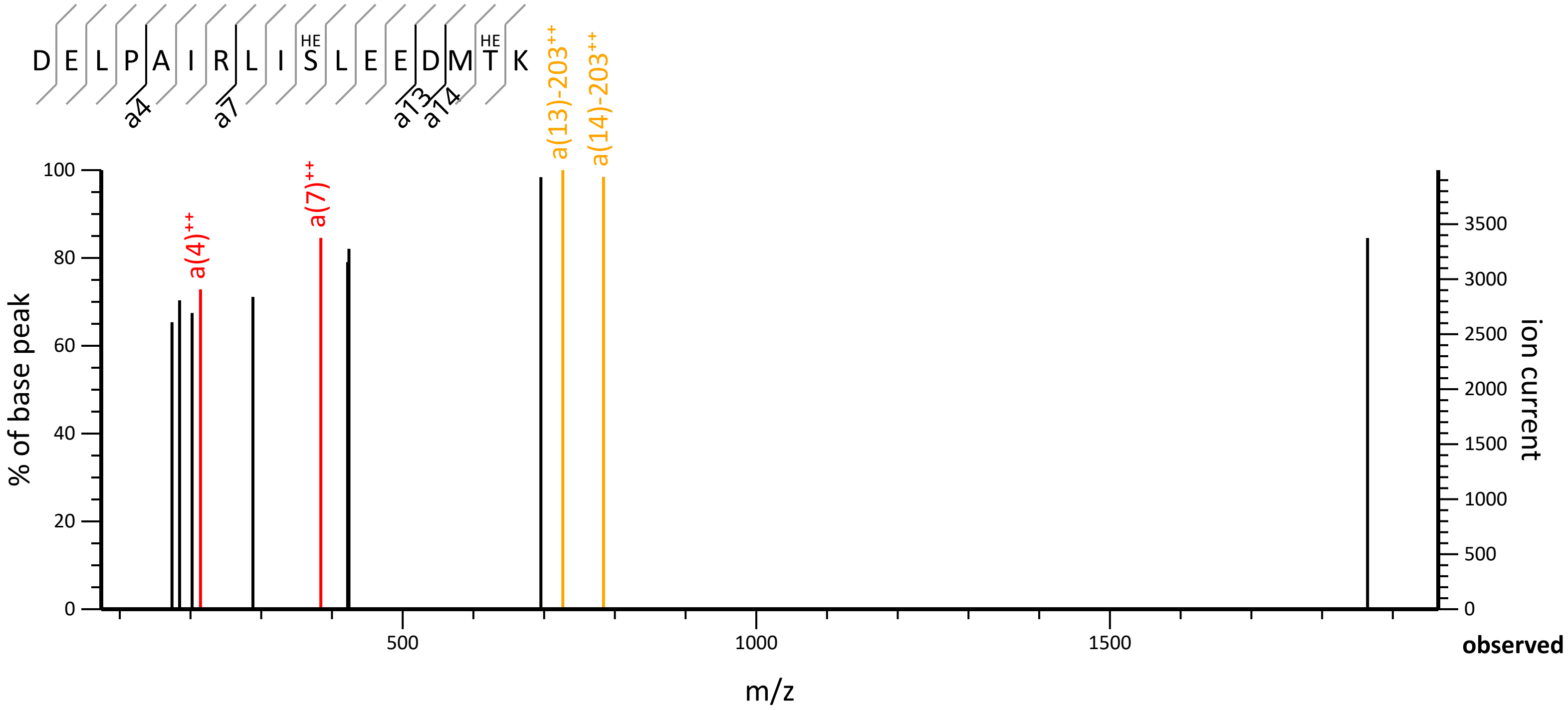

D E L P A I R L I S L E E D M T K
HE
HE
a4
a7
a13
a14
a(4)++
a(7)++
a(13)-203++
a(14)-203++
% of base peak
ion current
m/z
observed

Other BLAST web gateways

**All matches to this query**

| Score | Mr(calc) | Delta | Sequence |
|---|---|---|---|
| 22.2 | 2378.1774 | 2.0028 | DELPAIRLISLEEDMTK |
| 2.5 | 2379.1926 | 0.9875 | YTCGNRKVIPNMPDLILR |
| 0.0 | 2380.1456 | 0.0345 | METIWIPHLHTALAYMHER |
| 0.0 | 2378.2104 | 1.9697 | GSLEKLISESYKFIR |

**Mascot:** http://www.matrixscience.com/
